# Cancer-Associated Hypercalcemia Signals Through the Hindbrain to cause Anorexia

**DOI:** 10.1101/2024.03.19.585694

**Authors:** Diego Y. Grinman, Farzin M. Takyar, Pamela Dann, Marya Shanabrough, Stacey Brown, Alisson Clemenceau, Julie R. Hens, Bernardo Stutz, Lewis A. Chodosh, Wenhan Chang, Gerald I. Shulman, Tamas L. Horvath, John J. Wysolmerski

## Abstract

Hypercalcemia, caused by tumor secretion of parathyroid hormone-related protein (PTHrP), is associated with anorexia and weight loss. We demonstrate that overexpression of PTHrP by tumor cells in a transgenic model of breast cancer causes anorexia and rapid weight loss. These changes are accompanied by activation of neurons in the area postrema (AP), the nucleus tractus solitarius (NTS) and the parabrachial nucleus (PBN), a hindbrain circuit regulating food intake. Blocking hypercalcemia prevents anorexia and activation of these brain centers in tumor bearing mice, whereas injecting calcium activates the same circuit in wild-type mice. Neurons in the AP express the calcium-sensing receptor (CaSR) and the same AP/NTS/PBN circuit is stimulated by treating WT mice with cinacalcet, an allosteric activator of the CaSR. Finally, treating diet-induced obese mice with cinacalcet reduces food intake and causes weight loss. These results suggest that CaSR-expressing neurons in the AP might be a pharmacologic target for obesity.

## INTRODUCTION

Hypercalcemia is a frequent, and deadly complication of malignancy, occurring in up to 20%-30% of all cancer patients during the course of their disease. Cancer-associated hypercalcemia is typically observed in patients with advanced disease and often portends a poor prognosis, with a reported median survival between 25 to 52 days after initial elevation of calcium levels ^1–3^. Hypercalcemia can be the direct effect of osteolysis surrounding skeletal metastases or may result from a paraneoplastic syndrome known as humoral hypercalcemia of malignancy (HHM). HHM is most often caused by tumor cell secretion of parathyroid hormone-related protein (PTHrP) ^4^, a peptide growth factor related to parathyroid hormone (PTH) that, when present in the circulation, interacts with the shared PTH/PTHrP receptor (PTHR1) in kidney and bone to mimic the systemic actions of PTH ^5^.

Hypercalcemia frequently causes nausea, vomiting, anorexia and weight loss ^6^, symptoms that are also common in the cancer anorexia-cachexia syndrome (CACS). CACS is a condition of negative energy balance caused by tumor secretion of circulating factors that variably inhibit food intake, increase lipolysis, induce myolysis, and increase basal energy expenditure by browning of white adipose tissue (WAT). Different combinations of these effects occur in individual patients, but the end result is usually the depletion of peripheral glycogen, fat and protein stores, liberating glucose, lipids, and amino acids that fuel tumor cell growth. CACS is a lethal metabolic complication of cancer that contributes to 20% of all cancer deaths ^7^. Patients with HHM are frequently cachectic and it has been suggested that PTHrP contributes to CACS either by causing anorexia or by inducing the browning of WAT ^8–11^.

The calcium-sensing receptor (CaSR) is a G protein-coupled receptor (GPCR) most highly expressed on parathyroid and kidney cells, where it regulates PTH production and renal calcium excretion/reabsorption respectively ^12–14^. The CaSR is also expressed in other organs, including the brain. CaSR mRNA has been detected within neurons and glial cells in a variety of brain regions, including the area postrema and the hypothalamus, both of which regulate food intake^15–22^. However, CaSR functions in the brain are not well understood.

The area postrema (AP) lies outside the blood-brain barrier and acts as a chemosensory area, responding to a variety of circulating molecules associated with ingested toxins, systemic inflammation, cancer, or visceral malaise ^21,23–26^. Activation of AP neurons can induce food aversion, anorexia, nausea, and vomiting. AP neurons project to the adjacent nucleus tractus solitarius (NTS) and to the parabrachial nucleus (PBN). An important function of the PBN is to collect and integrate sensory input regarding noxious stimuli from a variety of peripheral and central centers and to relay “warning” signals to the forebrain ^21,23–26^. Activation of an AP/NTS/PBN pathway induces a powerful anorectic response leading to considerable weight loss in rodent models ^25^. Interestingly, this same pathway is activated by weight loss drugs such as glucagon-like peptide-1 receptor (GLP1R) agonists, gastric inhibitory polypeptide (GIP) receptor agonists, and amylin ^25,27–30^.

We produced a tetracycline-regulated transgenic model of PTHrP overexpression in mammary tumors to explore the metabolic consequences of PTHrP secretion in a mouse model of HHM. In this report, we demonstrate that PTHrP secretion from mammary carcinomas reproduces the cardinal features of HHM and causes severe and rapid weight loss. We found that weight loss was the result of anorexia and not the result of an increase in metabolic rate. However, we determined that anorexia is not a direct effect of PTHrP but, rather, the result of hypercalcemia activating a hindbrain AP/NTS/PBN circuit. We document CaSR expression within the AP, specifically by GLP1R-expressing neurons previously shown to be involved in food intake. Finally, we demonstrate that calcium, or the calcimimetic agent cinacalcet, reduces food intake and activates the AP/NTS/PBN pathway in WT mice, and that cinacalcet causes weight loss in mice with diet-induced obesity.

## METHODS

### Animals

All animals were housed in a temperature- and humidity-controlled room with a 12-h light/dark cycle. The animals were allowed free access to water and a standard laboratory diet, unless otherwise specified. All animal experiments were performed in accordance with institutional regulations after protocol review and approval by Yale University’s Institutional Animal Care and Use Committee.

We used previously described, MMTV-rtTA;TetO-PTHrP;MMTV-PyMT (Tet-PTHrP,PyMT) mice in a FVB background. In these mice, PTHrP expression can be turned on within PyMT-induced breast tumors by administering doxycycline (Dox) ^31^. Female Tet-PTHrP,PyMT mice with palpable tumors (about 10 weeks old) were treated with Dox (2mg/ml; Cat# D43020, Research Products International) in 5% sucrose water or with two subcutaneous pellets (15mg/pellet; Cat# B-168, Innovative Research of America) inserted on the lateral side of the neck with a trochar (Cat# MP-182, Innovative Research of America) and sealed with tissue adhesive (Cat# 1469, 3M). Five percent sucrose water or placebo pellets (15mg/pellet; Cat# C-111, Innovative Research of America) were used as controls. Four days after Dox administration, animals were euthanized, and tissue and blood were collected.

Male C57BL/6J mice were obtained from Charles River Laboratories and used in the high fat diet-induced obesity experiments (see below).

### Biochemical measurements

Plasma PTHrP was measured using an immunoradiometric assay (DSL-8100; Beckman Coulter) in which we substituted a rabbit anti-PTHrP (1–36) antibody generated in our laboratory as capture antibody ^31^. This assay has a sensitivity of 0.3 pM. Serum calcium concentrations were measured using the Quantichrom Calcium Assay Kit (DICA-500, BioAssay Systems) according to manufacturer’s instructions. Plasma carboxy-terminal telopeptide of type I collagen (CTX-1) levels were measured using an enzyme-linked immunosorbent assay (Cat AC-06F1, Ratlaps™) and procollagen type I collagen N-terminal propeptide (P1NP) with a competitive enzyme immunoassay (AC-33F1, IDS). PTH was determined in plasma samples using a two-site enzyme-linked immunosorbent assay (ELISA) (60-2305, Quidel). Plasma triglycerides (TG), non-esterified fatty acids (NEFA) and glycerol were determined using enzymatic kits (Triglyceride SL, Sekisui Diagnostics; NEFA-HR, Wako; glycerol, ab65337 Abcam) according to the manufacturer’s instructions. Serum Lipocalin-2 levels were measured by ELISA (DY1857, R&D Systems).

### Histology and immunohistochemistry

Gonadal adipose tissue was dissected, washed in phosphate-buffered saline (PBS) and fixed in Zinc-formalin (Z2902, Sigma) for 24h, dehydrated, and embedded in paraffin. Five μm sections were cut on a microtome and stained with hematoxylin and eosin (H&E) or used for immunohistochemistry. The later was performed using standard protocols. Briefly, deparaffinized sections were subjected to antigen retrieval using 10mM citrate buffer pH 6.0, after which endogenous peroxidase was quenched with 0.6% H2O2. Sections were washed with PBS and blocked with normal rabbit serum (PK-6105, Vector Labs) or 1% bovine serum albumin (BSA) in tris-buffered saline with 0.1% tween 20 (TBST) for 1h and then incubated overnight with Perilipin-1 (Ab61682, Abcam) and UCP-1 (Ab10983, Abcam) primary antibodies, respectively. After washing, sections were incubated with Biotinylated anti-goat (PK-6105, Vector Labs) or horseradish peroxidase (HRP) anti-rabbit (RMR 622-G, Biocare) and developed with 3,3’-diaminobenzidine (DAB, SK-4105, Vector Labs). Six pictures per slide were randomly acquired for the determination of adipocyte numbers and size in perilipin-1-immunostained and H&E-stained sections, and quantified using the ImageJ Adiposoft plugin ^32^.

For brain immunohistochemistry, mice were anesthetized and perfused via the left ventricle with heparinized saline [10,000 units/l heparin in 0.9% (w/v) NaCl] followed by 4% (w/v) paraformaldehyde plus 0.1% glutaraldehyde in phosphate buffer (PB) (0.1 M, pH 7.4). Brains were removed and post-fixed overnight at 4°C in glutaraldehyde-free fixative. Fifty μm vibratome sections were cut in the sagittal plane. After washing in PBS, sections were incubated at room temperature for 1h in blocking buffer [0.1 M phosphate buffer, 0.2% (v/v) Triton X-100, 10% (v/v) normal donkey serum (Cat# 017–000-121, Jackson Immunoresearch Labs). Sections were incubated in goat anti-c-Fos (1:5000, Biosensis) in 1% (v/v) blocking buffer overnight at room temperature. After washing with PB, sections were incubated in donkey anti-goat Alexa-Fluor 594 (1:500, Thermofisher) in PB for 1.5 hours at room temperature. Sections were mounted on slides using Vectashield fluorescence mounting media (Vector Labs) and images were visualized and captured using a Keyence BZ-X700 fluorescence microscope (Osaka, Japan). Quantification of c-Fos immunoreactivity was performed by counting the number of c-Fos-positive neurons on a region of interest using the “Analyze Particles” ImageJ plugin and represented as c-Fos density by dividing that number to the total ROI area.

Immunofluorescent staining of CaSR was detected using an AlexaFluor647-conjugated mouse monoclonal antibody (clone 3H8E9) raised against a peptide (*DYGRPGIEKFREEAEE*) in the extracellular domain of the CaSR at a concentration of 5 μg/ml and counterstained with SYTO13 fluorescent nucleic acid dye (ThermoFisher, S7575) as previously described ^33^. Fluorescent images of whole brain sections were acquired with a 20x objective and exported as dual-channel TIFF files from a GeoMx Digital Spatial Profiler instrument (NanoString Technologies) and presented in digitally assigned pseudo colors (CaSR in red and nucleus in blue).

### Fluorescent in situ hybridization

Fluorescent *in situ* hybridization (ISH) assays were performed in formalin-fixed, paraffin-embedded (FFPE) 5 μm sagittal brain sections using the RNAscope™ Multiplex Fluorescent V2 system (Advanced Cell Diagnostics) to visualize RNA transcripts in the AP, NTS & PBN. Probes purchased from Advanced Cell Diagnostics included *Cfos* (316921), *Casr* (423451), *Glp1r* (418851) and *Cgrp* (578771). Fixation, dehydration, hybridization, and staining protocols for FFPE sections were performed according to the manufacturer’s specifications. Negative and positive control images for each channel (negative control probe: *DapB*, positive control probes: *Polr2a* in channel C1, *Ppib* in channel C2, *Ubc* in channel C3 and *Hprt* in channel C4) were used to determine brightness and contrast parameters for experimental images. Images were acquired using a Leica TCS SP8 STED Confocal Super-Resolution Microscope with a frame size of 1,024 × 1,024 pixels using tiling mode and a 40× oil-immersion objective. Three brain slices from each animal were analyzed. Cells were counted manually with the cell counter plugin of ImageJ. *Cfos*-, *Casr*-, *Cgrp*- and *Glp1r*-positive cells were quantified in their corresponding separate channels using DAPI-labeled nuclei to identify individual cells. The respective different channels were overlapped to identify cells with positive signals for more than one probe.

### RNA extraction and real time RT-PCR

Total RNA from gonadal and brown interscapular adipose tissue, hypothalamus and AP lysates was isolated using PureLink RNA columns (Cat# 12183025, Invitrogen) following the manufacturer’s instructions. For all samples, the ratio of absorbance at 260 nm to absorbance at 280 nm was >1.8. For quantitative real-time PCR analysis, cDNA was synthesized using a maximum of 1 μg of total RNA with the High-Capacity cDNA Reverse Transcription Kit (Cat# 4368814, Applied Biosystems). Quantitative RT-PCR was performed using the Taqman Fast Universal PCR Master Mix (Cat# 4352042, Applied Biosystems) and a StepOnePlus real-time PCR system (Applied Biosystems). The following TaqMan primer sets were used: *Ucp1* (Mm01244861_m1), *Dio2* (Mm00515664_m1), *Ppargc1* (Mm01208835_m1), *Crh* (Mm01293920_s1), *Oxt* (Mm01329577_g1), *Trh* (Mm01182425_g1), *Pmch* (Mm01242886_g1), *Orx* (Mm01964030_s1), *Cartpt* (Mm04210469_m1), *Pomc* (Mm00435874_m1), *Npy* (Mm00445771_m1) and *Agrp* (Mm00475829_g1). *BActin* (Cat# 4352933E) and *Hprt* (Mm03024075_m1) were used as housekeeping. Relative mRNA expression was determined using the standard curve method with the StepOne software v2.3 (Applied Biosystems).

### Adipocyte isolation and treatment

Adipocytes were isolated by collagenase digestion of subcutaneous (SQ) fat pads of adult female FVB mice. Animals were euthanized and bilateral #4 and #5 SQ fat pads were dissected, the lymph node was removed, and the remaining fat pad was rinsed with PBS. Adipose tissue was minced into 3-4mm pieces and transferred to 50 ml conical tubes containing 10 ml of digestion buffer [Kreb-Ringers-HEPES buffer pH 7.4 (10 mM HEPES, 2.2 mM CaCl2, 120 mM NaCl, 1.2 mM KH2PO4, 1.2 mM MgSO4, 4.7 mM KCl) + 1 mg/ml type VIII collagenase (C2139, Sigma)], mixed by inversion and incubated at 37 °C for 1h in a shaking water bath at 150 rpm. The dissociated mixture was then filtered using a 230 µm filter and centrifuged for 4 min at 200xg to float adipocytes and pellet the stromal vascular fraction (SVF). The SVF and infranatant were then removed with a glass Pasteur pipette and the remaining adipocyte-enriched fraction was washed with working buffer (KRH + 2.5 mM glucose + 1% Fatty acid free BSA + 200 nM adenosine). The adipocytes were gently aspirated, divided into separate tubes containing 5 ml of the working buffer and either vehicle, isoproterenol 20 µM or PTHrP 1-34 10 nM (4017147, Bachem) and incubated for 30 min at 37 °C under agitation. At this point, the adipocytes were centrifuged for 4 min 200xg, 500 ul of the infranatant was saved for free glycerol analysis (see below) and the rest, along with the pellet, discarded. The remaining layer of floating adipocytes was lysed for protein analysis using RIPA buffer supplemented with a cocktail of protease inhibitors (Thermo Scientific Cat#78429), 50 mM NaF, and 1 mM Na3VO4 on ice, and cytosolic and fat-associated protein extract were obtained as previously reported ^34^.

Free glycerol was measured in the infranatant fraction of the isolated adipocyte culture using a colorimetric assay kit (ab65337, Abcam) following the manufacturer’s instructions.

### Western blot

Western blot analysis was conducted as previously described ^31^. Briefly, total protein was measured in samples using the Bradford protein assay (Bio-Rad Cat# 5000001) following the manufacturer’s instructions. A 2µg/µl protein solution containing sample buffer (Invitrogen Cat# NP0007) plus sample reducing agent (Invitrogen Cat# NP0004) was prepared and 30ug of total protein were loaded into precast, 4% to 12% Bis-Tris acrylamide gels (Thermo Fisher Scientific, Cat# NP0322) in MOPS buffer (Thermo Fisher Scientific, Cat# NP0001). Following electrophoresis, samples were transferred to nitrocellulose membranes (Bio-Rad, Cat# 1621112). Membranes were treated with blocking buffer (LI-COR Biosciences, Cat# 927-60001) for 1 hour at room temperature and then incubated with the primary antibody overnight at 4°C, followed by a dye conjugated secondary antibody for 1 hour at room temperature. Membranes were imaged and analyzed using the Odyssey IR imaging system (LI-COR Biosciences). The primary antibodies used were: pSer(563)HSL (Cat# 4139), pSer(660)HSL (Cat# 4126), tHSL (Cat# 4107) from Cell signaling, βActin (Cat# sc-130656) from Santa Cruz Biotechnology and perilipin-1 (Cat# Ab61682) from Abcam. The secondary antibodies used were anti-mouse (Cat# 926-68022), anti-Rabbit (Cat# 926-32213) and anti-goat (Cat# 926-32214) from LI-COR Biosciences.

### Body composition measurements and metabolic cage studies

Energy expenditure and caloric intake were measured in a Comprehensive Laboratory Animal Monitoring System (CLAMS; Columbus Instruments). Tet-PTHrP mice and WT control were study sequentially, first while ingesting 5% sucrose water (Off Dox) for 4 days and then while ingesting 5% sucrose water containing 2mg/ml of Dox (On Dox) for another 4 days. Mice were allowed to acclimate to the metabolic cages for 24 hours before beginning data collection on each interval. Body composition was measured by ^1^H magnetic resonance spectroscopy using a Bruker Minispec analyzer.

### Animal Treatments

10 mg/kg of Osteoprotegerin (OPG-Fc, Amgen), 5 mg/kg of zoledronic acid (ZA; SML0223, Sigma), 75ug/mouse of anti-LCN2 (MAB1857, R&D systems) & 10 mg/kg of anti-PTHR1 antibody (XOMA) & control IgG, were subcutaneously administrated to female Tet-PTHrP,PyMT mice with palpable tumors (about 10 weeks old) simultaneously with Dox pellet implantation.

#### Calcium gluconate and cinacalcet treatment

10-week-old female mice were placed in individual cages, fasted overnight and then injected intraperitoneally (i.p.) with one dose of 1 mmol/kg calcium gluconate or equal volumes of vehicle (saline) every 30 min. Other mice were injected with a single dose of 30 µmol/kg cinacalcet or equal volumes of vehicle [20% (2-Hydroxypropyl)-β-cyclodextrin (HβC)]. Food was added to the cage at the time of the first injection and food consumption during the next 2 hours was assessed by comparing the weight of the food added with that left over at the end of the time-period. At the end of the 2 hours, mice were anesthetized with isofluorane, blood was collected by cardiac puncture and then mice were sacrificed, and brains harvested as noted above.

### Diet induced obesity

Male C57BL/6J litters were fed with a 45 % (Kcal) fat diet (Research Diets, Cat# D12451) for 12 weeks starting at the time of weaning. Animals weighing ≥30g ^35^ were individually caged and baseline daily food intake and weight were monitored for 7 days using a digital scale. Subsequently, obese mice were randomly assigned to receive 7 days of: 1) daily i.p. injections of 15 µmol/kg cinacalcet in HβC; or 2) daily i.p. injections of HβC alone. Food intake and weight were monitored daily until the conclusion of the experiment.

### Pair feeding

Tet-PTHrP;PyMT mice with tumors were treated with SQ Dox pellets for 4 days and the same amount of food consumed by this group was provided to mice treated with a SQ placebo pellets for each matching 24-hour period. A third group of placebo treated Tet-PTHrP;PyMT mice was also fed ad libitum and used as controls. Food intake and weight were measured daily for each animal. Body fat, lean and free fluid composition was determined by ^1^H magnetic resonance spectroscopy at the end of the experiment.

### Statistics

Results were expressed as mean ± standard error of the mean (SEM) of at least 3 independent experiments. Statistical analyses were performed with Prism 9.0 (GraphPad Software) and consisted of Student’s t-tests (two-sided with 95% confidence level) in the statistical analysis of two groups, one-way ANOVA followed by Tukey’s or Dunnet’s multiple comparisons tests for more than two groups with one variable and Pearson’s correlation analysis to examine the relationship between circulating calcium levels and food intake. Before statistical analysis, Q-Q plot and Shapiro Wilks test were performed for normality. Homoscedasticity was assessed with Levene’s test. In figures, asterisks mean significant differences between means. Differences were considered significant at p < 0.05.

## RESULTS

### Tumor secretion of PTHrP causes hypercalcemia and weight loss

In order to investigate the metabolic consequences of HHM, we created a tetracycline-regulated model of PTHrP overexpression in transgenic mammary tumors ^31^. We generated triple transgenic, MMTV-rtTA;pTet-PTHrP;MMTV-PyMT (Tet-PTHrP;PyMT) mice in which the reverse tetracycline transactivator (rtTA) and the PyMT oncogene are both expressed in mammary epithelial cells under control of the mouse mammary tumor virus long terminal repeat (MMTV). As a result, administration of Dox to Tet-PTHrP;PyMT mice activates human *PTHLH* cDNA expression in mammary tumors.

We treated female, Tet-PTHrP;PyMT mice with Dox after they had developed palpable tumors (approximately 10 weeks old). Tumor-bearing, control MMTV-PyMT mice and tumor-bearing, control Tet-PTHrP;PyMT mice off Dox had low levels of circulating PTHrP and normal circulating calcium levels. However, tumor bearing, Tet-PTHrP;PyMT mice treated with Dox for 4 days demonstrated elevated circulating levels of PTHrP and severe hypercalcemia ^31^ (Fig. 1A). Dox treatment also increased circulating CTX-1, a marker of bone resorption, without significantly changing circulating P1NP, a marker of bone formation, consistent with the known effects of PTHrP on skeletal turnover ^5^ (Extended data Fig. 1A). In addition to hypercalcemia, overexpression of PTHrP in Tet-PTHrP;PyMT mice on Dox was also associated with a 17% reduction in body weight over 4 days of Dox treatment (Fig. 1B). At sacrifice, Tet-PTHrP;PyMT mice on Dox showed severe atrophy of the gonadal, mesenteric, retroperitoneal and interscapular adipose tissue depots as compared to controls (Fig. 1C).

**Figure 1.**
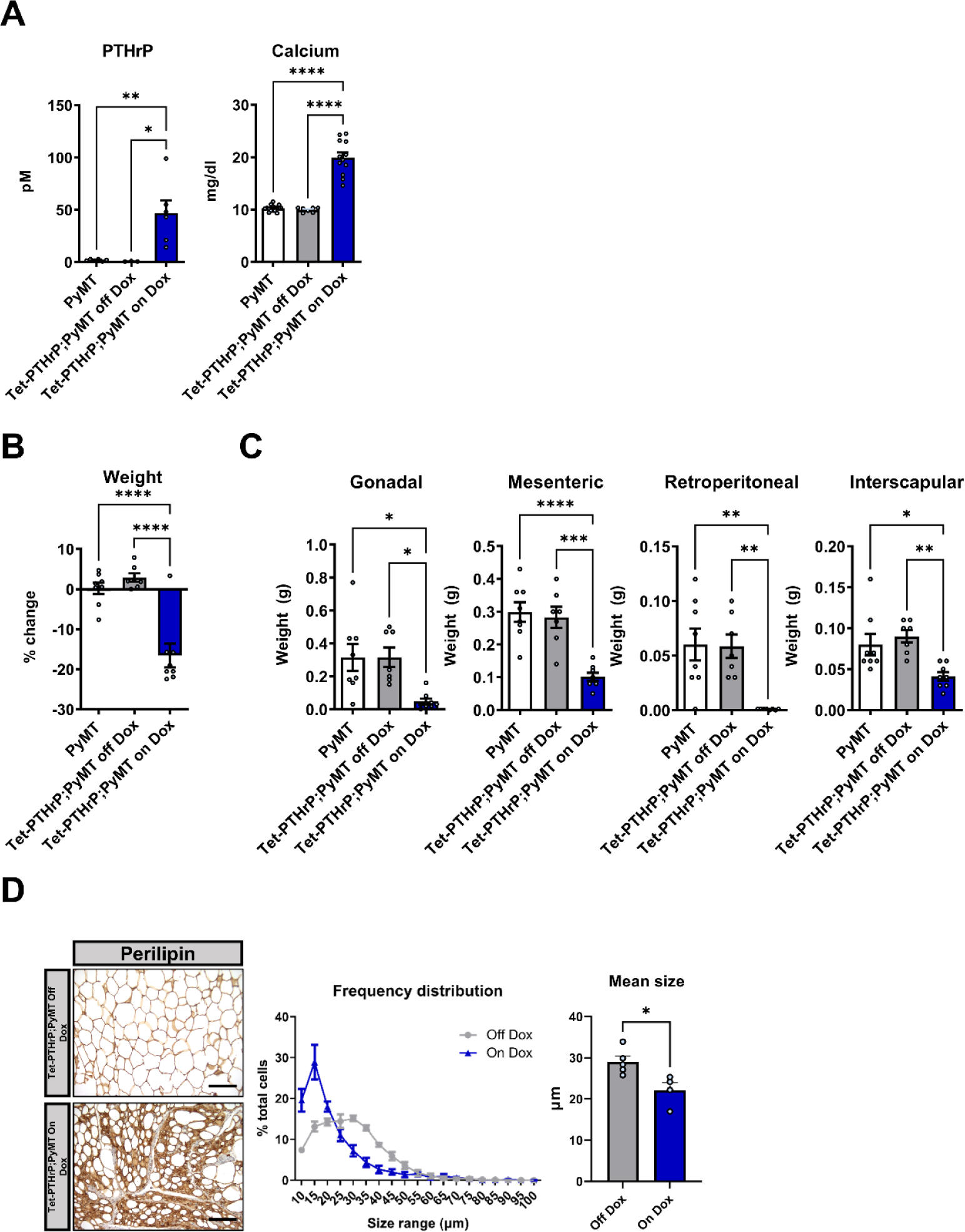
Tumor PTHrP production causes weight loss. 10-week-old female Tet-PTHrP;PyMT mice with palpable tumors were treated with Dox for 4 days; PyMT mice and Tet-PTHrP;PyMT mice were treated with 5% sucrose water as controls. (A) Circulating levels of plasma PTHrP and serum calcium concentration. (B) Percent change in body weight over the course of the experiment. (C) Weights of different fat depots at time of sacrifice. (D) Representative images of perilipin immunostaining of gonadal WAT from Tet-PTHrP;PyMT Off Dox and On Dox, and histomorphometric quantification of adipocyte size. Scale bars: 100µm. Bars in each graph represent mean ± SEM, with a minimum of n=4 mice. ****p<0.0001 ***p<0.001 **p<0.01 *p<0.05.

### Overexpression of PTHrP in Mammary Tumors Induces Lipolysis by Stimulating the PTHR1

Histological examination of the gonadal WAT (gWAT) suggested that PTHrP overexpression was associated with a reduction in adipocyte size (Extended data Fig. 1B). A more formal histomorphometric analysis of perilipin-stained gWAT confirmed this impression. As shown in Fig. 1D, mice overexpressing PTHrP had significantly smaller adipocytes as defined by both a reduction in mean adipocyte size as well as a shift in the size distribution of adipocytes towards smaller cells in gWAT from Tet-PTHrP;PyMT mice on Dox vs. off Dox. There was also an increase in circulating levels of glycerol, triglycerides, and non-esterified free fatty acids (NEFA) upon Dox treatment, although the changes in NEFA did not reach statistical significance (p=0.07) (Extended data Fig. 1C).

The effects of PTHrP on bone resorption and circulating calcium levels are known to be mediated by the PTHR1, although it is not known whether the same receptor would mediate weight loss and fat wasting. Therefore, we treated Tet-PTHrP;PyMT mice on Dox with either an anti-PTHR1 blocking antibody or IgG Isotype control ^31^. As expected, blocking the PTHR1 completely normalized calcium levels, reduced CTX-1 levels and increased P1NP levels, without significantly altering PTHrP levels (Extended data Fig. 2A&B). Administration of the anti-PTHR1 antibody also prevented weight loss and was associated with a significant inhibition of fat loss as reflected by the increased gonadal, mesenteric and interscapular adipose tissue weights in mice treated with the anti-PTHR1 compared to IgG controls (Extended data Fig. 2C&D). Treatment with the anti-PTHR1 also resulted in the maintenance of mean adipocyte sizes in the face of PTHrP overexpression, suggesting that the reduction in adipocyte size depended on PTHR1 signaling. Finally, blocking the PTHR1 reversed the elevations in circulating triglycerides, glycerol and NEFA that were associated with mammary tumor PTHRP overexpression (Extended data Fig. 1C). All together, these findings suggested that overexpression of PTHrP in tumor-bearing mice caused increased WAT lipolysis by acting on the PTHR1.

**Figure 2.**
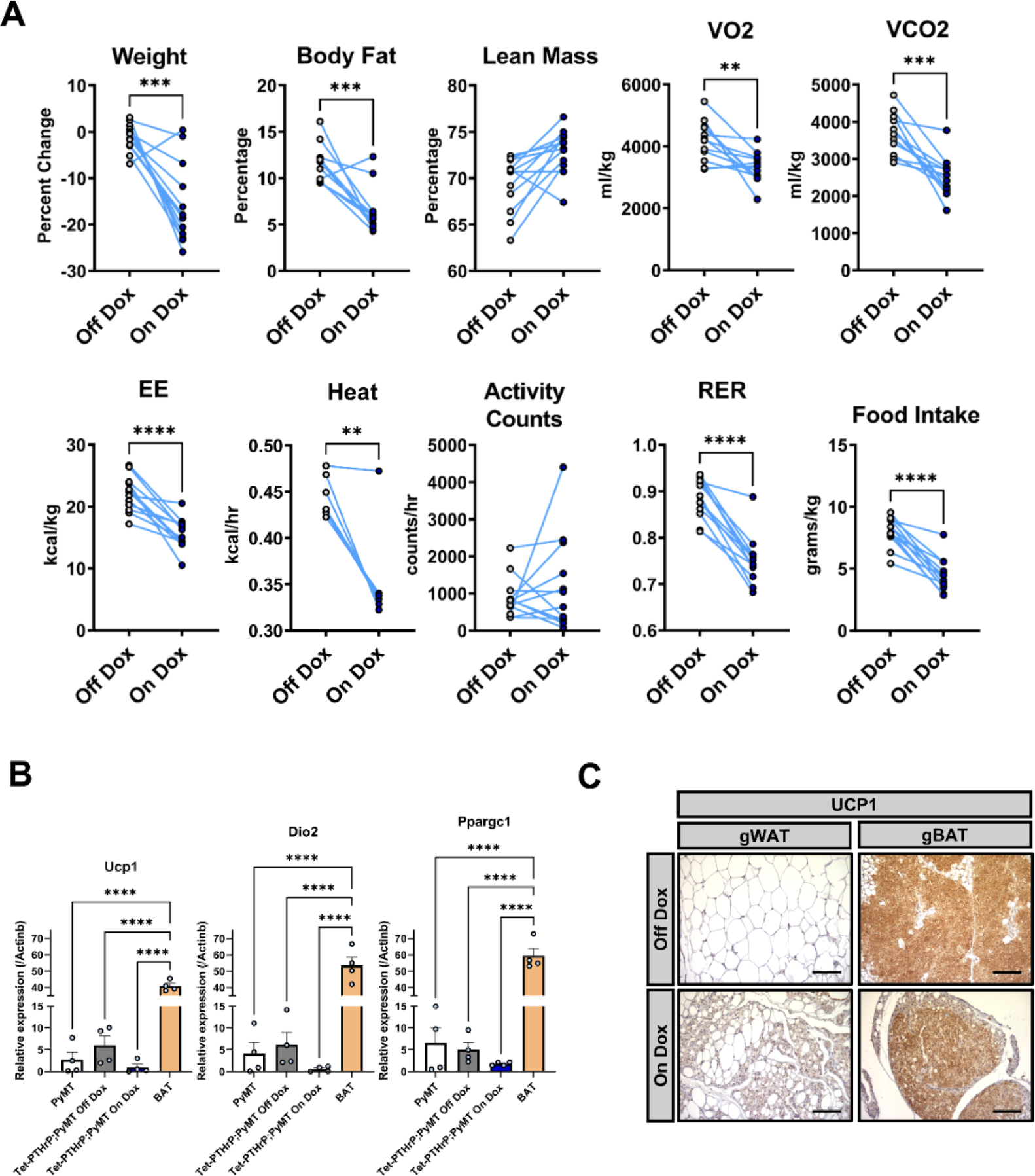
Overexpression of PTHrP reduces food intake and lowers metabolic rate. A) Female Tet-PTHrP;PyMT mice were placed in metabolic cages and were studied during 2 successive intervals: first on 5% sucrose water for 5 days and then on Dox for another 5 days. Mice were weighed and body fat and lean mass were measured by ^1^H-NMR before and after each experimental period. Dots connected by a line represents data from one mouse during time periods off Dox (left dot) and on Dox (right dot); n=12. Each graph represents a different metabolic measurement as noted at the top of the graph. Abbreviations: VO2, O2 consumption; VCO2, CO2 production; EE, energy expenditure; RER, respiratory exchange ratio. B) QPCR analysis of gonadal white adipose tissue (gWAT) harvested from PyMT mice, Tet-PTHrP;PyMT mice off Dox and Tet-PTHrP;PyMT mice on Dox in order to measure the expression of thermogenic genes serving as markers of brown adipose tissue. Interscapular brown adipose tissue (BAT) was used as positive control. *βActin* was used as a housekeeping gene to control for variations in RNA content. Bars represent mean ± SEM, n=4. C) Representative images of UCP1 immunostaining of gWAT and gBAT from Tet-PTHrP;PyMT Off or On Dox. Scale bars: 100µm. ****p<0.0001 ***p<0.001 **p<0.01.

PTH and PTHrP have previously been shown to induce lipolysis in primary cultures of white adipocytes through activation of the PTHR1 ^36^. Therefore, we next cultured freshly isolated adipocytes from WT FVB mice with PTHrP for 30 min to confirm that PTHrP could act directly on adipocytes to cause lipolysis. PTHrP treatment activated hormone sensitive lipase (HSL) in these cells as evidenced by HSL phosphorylation at 2 canonical sites, Ser563 and Ser660. PTHrP also stimulated glycerol release into cultured media (Extended data Fig. 1D). These findings support the fact that PTHrP secretion from mammary tumors could act systemically to increase adipocyte lipolysis.

### Overexpression of PTHrP in Mammary Tumors Causes Anorexia and Reduced Energy Expenditure

Previous studies suggested that PTHrP caused weight loss, fat wasting and muscle atrophy due to “browning” of WAT and a hypermetabolic state ^8,37^. Therefore, we assessed systemic energy metabolism in Tet-PTHrP;PyMT mice using metabolic cages to measure physical activity, food intake and energy expenditure by indirect calorimetry (Fig. 2A). Tet-PTHrP;PyMT mice were studied sequentially, first off Dox and then on Dox, allowing each mouse to serve as its own control. We also measured body composition using proton NMR at the beginning and end of each metabolic cage session. As noted previously, PTHrP expression in mammary tumors caused weight loss and a significant reduction in total body fat by proton NMR. However, rather than increasing metabolic rates, tumor PTHrP overexpression reduced oxygen consumption (V_O2_), carbon dioxide production (V_CO2_), overall energy expenditure (EE), and heat production without significantly changing activity. PTHrP overexpression also significantly decreased the respiratory exchange ratio (RER) (CO_2_ produced in proportion to O_2_ consumed), suggesting increased fatty acid oxidation. Finally, food consumption was reduced by 44% on Dox treatment. WT mice treated with Dox for 4 days had no alterations in body weight, fat content or energy expenditure, ruling out independent effects of Dox treatment (Extended Fig. 3).

**Figure 3.**
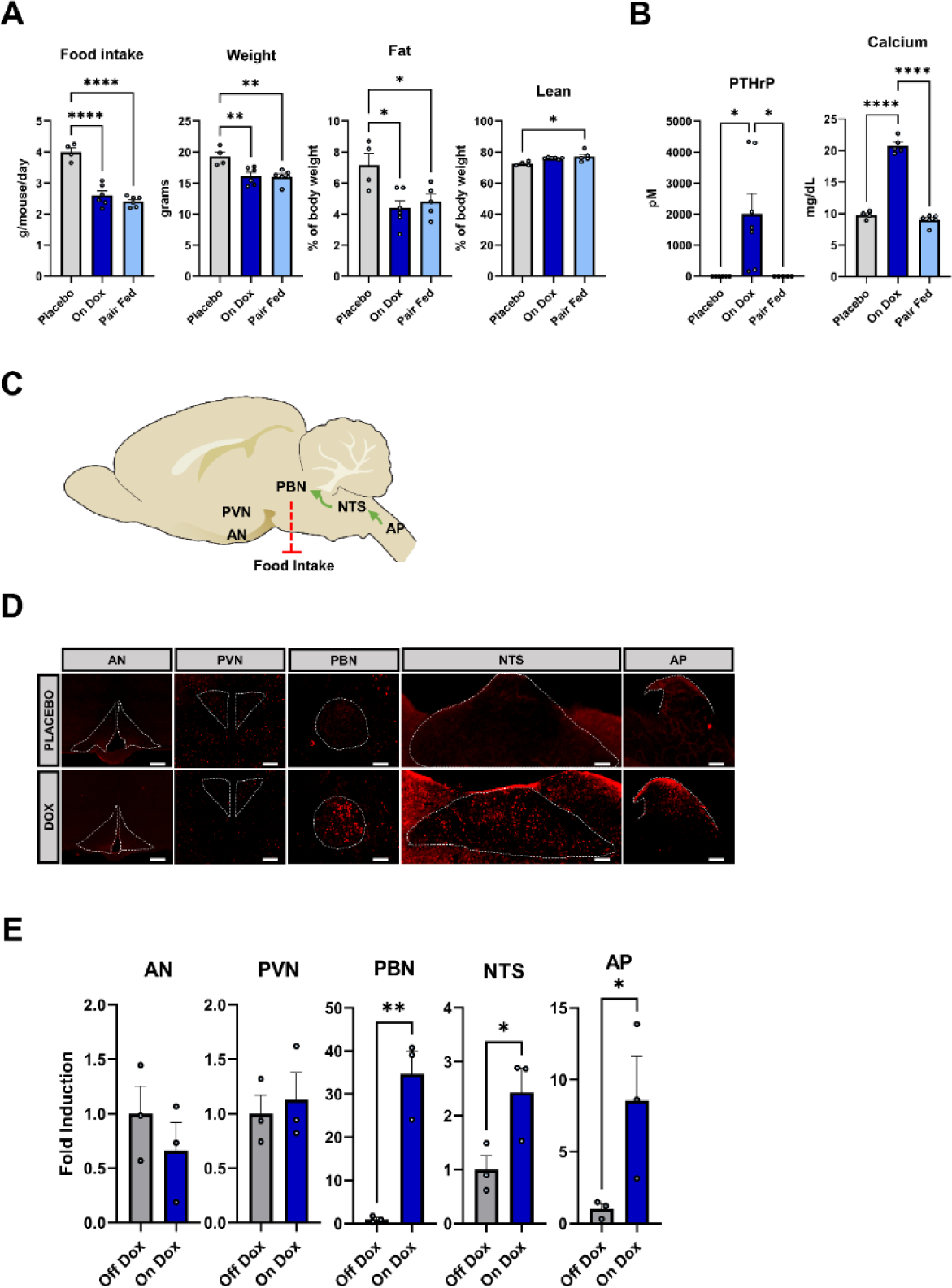
Anorexia caused by tumor PTHrP production is associated with neuronal activation in the AP, NTS and PBN. A) Food intake, body weight and body composition in control and pair-fed mice. Daily food intake was measured in Tet-PTHrP;PyMT mice treated with subcutaneous Dox pellets (on Dox) and the same amount of food was given to Tet-PTHrP;PyMT mice on placebo pellets (Pair Fed) for 4 days. Tet-PTHrP;PyMT mice on placebo fed *ad libitum* (Placebo) served as controls. Weight refers to body dry weight, which was calculated by subtracting the free fluid mass from the whole-body weight. Body fat % and lean % composition was determined by ^1^H NMR. A minimum of 4 mice were studied in each group. B) Plasma PTHrP concentrations and serum calcium concentrations in the three groups of mice (Placebo, On Dox, and Pair Fed) as described in A. C) Schematic representation of the mouse brain in a sagittal plane showing the Area Postrema (AP), Nucleus Tractus Solitarius (NTS) and Parabrachial Nucleus (PBN) anorectic circuit as well as the Arcuate Nucleus (AN) and Paraventricular Nucleus (PVN) ^68^. D) Representative images of c-Fos (red fluorescence) immunostaining of vibratome sections of brains in the coronal (AN & PVN) and sagittal (AP, NTS and PBN) plane from Tet-PTHrP;PyMT mice treated with Placebo or Dox. E) Bar graphs showing the quantification of c-Fos positive cells per unit area in 3 separate mice for each condition. Data are shown as fold induction over baseline off Dox. Scale bars: 100µm. Bars in all graphs represent mean ± SEM, ****p<0.0001 **p<0.01 *p<0.05.

We also evaluated whether PTHrP overexpression caused white adipose tissue browning by using QPCR to compare the expression of brown adipose tissue (BAT) marker mRNAs in gWAT from Tet-PTHrP;PyMT mice on Dox as compared to controls (Fig. 2B). Overexpression of PTHrP did not change the expression of *uncoupling protein 1* (*Ucp-1*), *iodothyronine deiodinase 2* (*Dio2*) or *peroxisome proliferator-activated receptor-gamma coactivator 1 alpha* (*PGC-1α*). In addition, as noted before, although gWAT from Tet-PTHrP;PyMT mice on Dox showed a marked reduction in adipocyte size (Fig. 1D & Extended Fig. 1B), no multilocular adipocytes were evident. Furthermore, immunostaining of gonadal fat from Tet-PTHrP;PyMT mice off and on Dox revealed clusters of BAT with high expression of UCP1 as previously reported ^38^. However, we failed to detect UCP1 in the WAT compartment in either group (Fig. 2C). Overall, these results suggest that, in our model, tumor PTHrP secretion does not cause browning of WAT or increased energy expenditure. Rather, it results in anorexia, lipolysis, increased fatty acid fuel utilization and reduced overall whole-body energy expenditure.

### Reduced Food Intake Reproduces Weight Loss Caused by PTHrP Overexpression

To evaluate whether the decrease in fat content and body weight induced by tumor PTHrP production were the consequence of lipolysis, reduced food intake, or both, we performed a pair feeding experiment. We matched the daily food intake of Tet-PTHrP;PyMT mice treated with placebo to that of Tet-PTHrP;PyMT mice treated with Dox for 4 days. Pair feeding resulted in equivalent reductions in body weight and percent body fat between the two groups of Tet-PTHrP;PyMT mice either treated with Dox and having elevated PTHrP and calcium levels or treated with placebo and having normal PTHrP and calcium levels (Fig. 3A&B). These results demonstrate that reduced caloric intake in Tet-PTHrP mice on Dox was the dominant factor driving weight loss and reductions in body fat.

The prominent anorexia associated with overexpression of PTHrP in tumors from Tet-PTHrP;PyMT mice focused our attention on brain centers known to regulate food intake. The arcuate nucleus (AN) and the paraventricular nucleus of the hypothalamus (PVN) are well described as central nervous system centers regulating appetite ^39^. Likewise, brainstem neurons in the AP and NTS, that project to the PBN also play a key role in regulating feeding behavior ^40^. We explored whether tumor PTHrP production stimulated neurons in either of these important appetite centers by performing immunofluorescence staining for c-Fos, a standard indicator of neuronal activation ^41^. As shown in Fig. 3D&E, we saw no c-Fos-positive neurons in the AN or in the PVN of anorectic, Tet-PTHrP;PyMT mice on Dox or in controls off Dox. Consistent with the lack of c-Fos staining in the hypothalamus, overexpression of PTHrP in mammary tumors produced no significant alterations in the expression of mRNAs for typical orexigenic (AgRP, NPY, ORX, MCH) or anorexigenic factors (POMC, TRH, OXY) within the hypothalamus (Extended data Fig. 4). There was an increase in the levels of CRH expression and a slight decrease in the CART mRNA levels upon Dox treatment, however neither of these transcripts were significantly different than the pair fed controls. As expected, the mRNAs for orexigenic peptides NPY and AGRP were increased in pair-fed, caloric-restricted control mice. However, it is significant that the same orexigenic mRNAs were not induced in Tet-PTHrP;PyMT mice on Dox despite their weight loss. In contrast to these results, tumor PTHrP secretion resulted in strong nuclear c-Fos staining of neurons within the AP, NTS and PBN, which was not present in control Tet-PTHrP;PyMT mice off Dox. Therefore, PTHrP production by mammary tumors appears to cause anorexia by activating a well-described AP/NTS/PBN pathway within the hindbrain.

**Figure 4.**
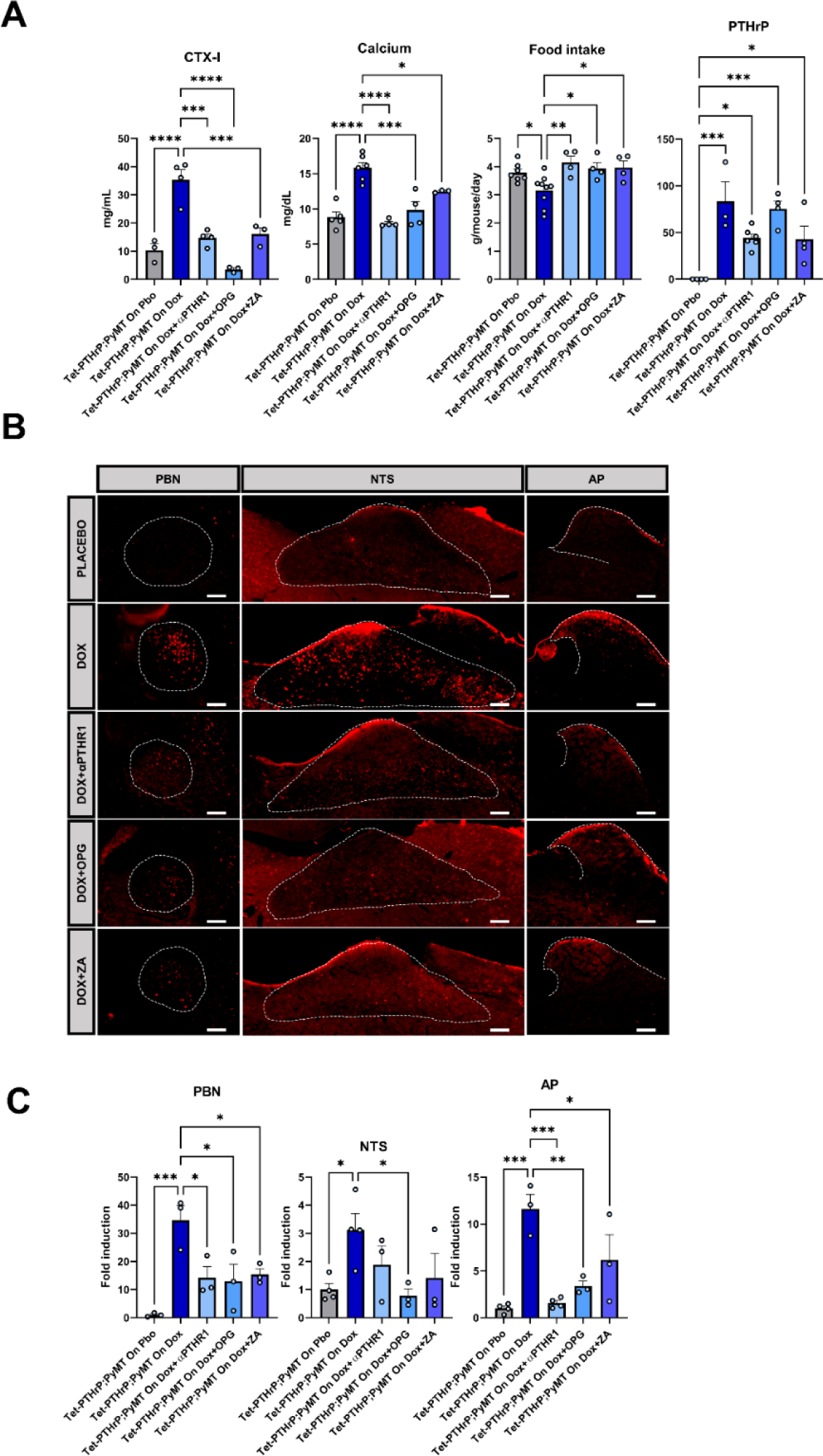
Preventing hypercalcemia limits anorexia and AP/NTS/PBN activation in tumor bearing mice. Tet-PTHrP;PyMT mice with tumors were treated for 4 days with Placebo, Dox, or Dox plus 10mg/kg of the anti-PTHR1 antibody (αPTHR1), Dox plus 10mg/kg of Osteoprotegerin (OPG) or Dox plus 5mg/kg of zoledronic acid (ZA). A) Circulating levels of plasma CTX-I, serum calcium concentration, daily food intake and circulating levels of PTHrP for the 5 different groups of mice. B) Representative images of c-Fos (red immunofluorescence) immunostaining of vibratome sections of the AP, NTS and PBN in the sagittal plane. C) Quantification of c-Fos positive cells per unit area in the 4 treatment conditions relative to control Tet-PTHrP;PyMT mice on Placebo. Scale bar: 100µm. Bars in each graph represent mean ± SEM, n=3 to n=9 mice per group. ****p<0.0001 ***p<0.001 **p<0.01 *p<0.05.

### Calcium, not PTHrP, activates the AP/NTS/PBN circuit and inhibits food intake

Tumor PTHrP overexpression results in high levels of circulating PTHrP that stimulate bone resorption and cause hypercalcemia. Furthermore, as outlined above, blocking the PTHR1 prevents weight loss but also blocks bone resorption and normalizes calcium (see Extended data Fig. 2A-C). To distinguish whether elevations in circulating PTHrP or elevated plasma calcium levels were responsible for reduced food intake, we treated Tet-PTHrP;PyMT mice on Dox with two different drugs, (Osteoprotegerin, (OPG-Fc) and zoledronic acid (ZA)), both of which inhibit bone resorption independent of effects on PTHrP or the PTHR1. As controls, we used Tet-PTHrP;PyMT mice on Dox with or without anti-PTHR1 antibody treatment. As expected, Tet-PTHrP;PyMT mice had elevated CTX-1 levels, were hypercalcemic, had reduced food intake and demonstrated c-Fos staining in the AP, NTS and PBN (Fig. 4A-C). Treatment with the anti-PTHR1 antibody blocked all these effects of PTHrP overexpression by tumors. Significantly, both OPG-Fc and ZA also lowered CTX-1 levels, reduced circulating calcium levels, prevented any reduction in food intake, despite persistent increases in circulating PTHrP levels. As with the anti-PTHR1 antibody treatment, both OPG-Fc and ZA also blocked the ability of tumor PTHrP production to activate neurons in the AP/NTS/PBN pathway (Fig. 4A-C). Collectively, these results suggest that bone resorption and/or hypercalcemia, rather than PTHrP, is required for activation of the anorexigenic AP/NTS/PBN circuit.

Lipocalin-2 (LCN2), also known as neutrophil gelatinase-associated lipocalin (NGAL), has been previously shown to be released from bone and to inhibit appetite in mice and in primates ^42,43^. Therefore, we measured LCN2 and found that overexpression of PTHrP in mammary tumors was associated with an increase in circulating LCN2 (Extended data Fig. 5A). However, treating Tet-PTHrP;PyMT mice on Dox with an anti-LCN2 blocking antibody did not result in any changes in food intake (Extended data Fig. 5B) or in c-Fos expression in the AP/NTS/PBN (Extended data Fig. 5C) as compared to control Tet-PTHrP;PyMT mice on Dox treated with vehicle. These results demonstrate that LCN2 does not mediate the anorectic response to tumor production of PTHrP and hypercalcemia.

**Figure 5.**
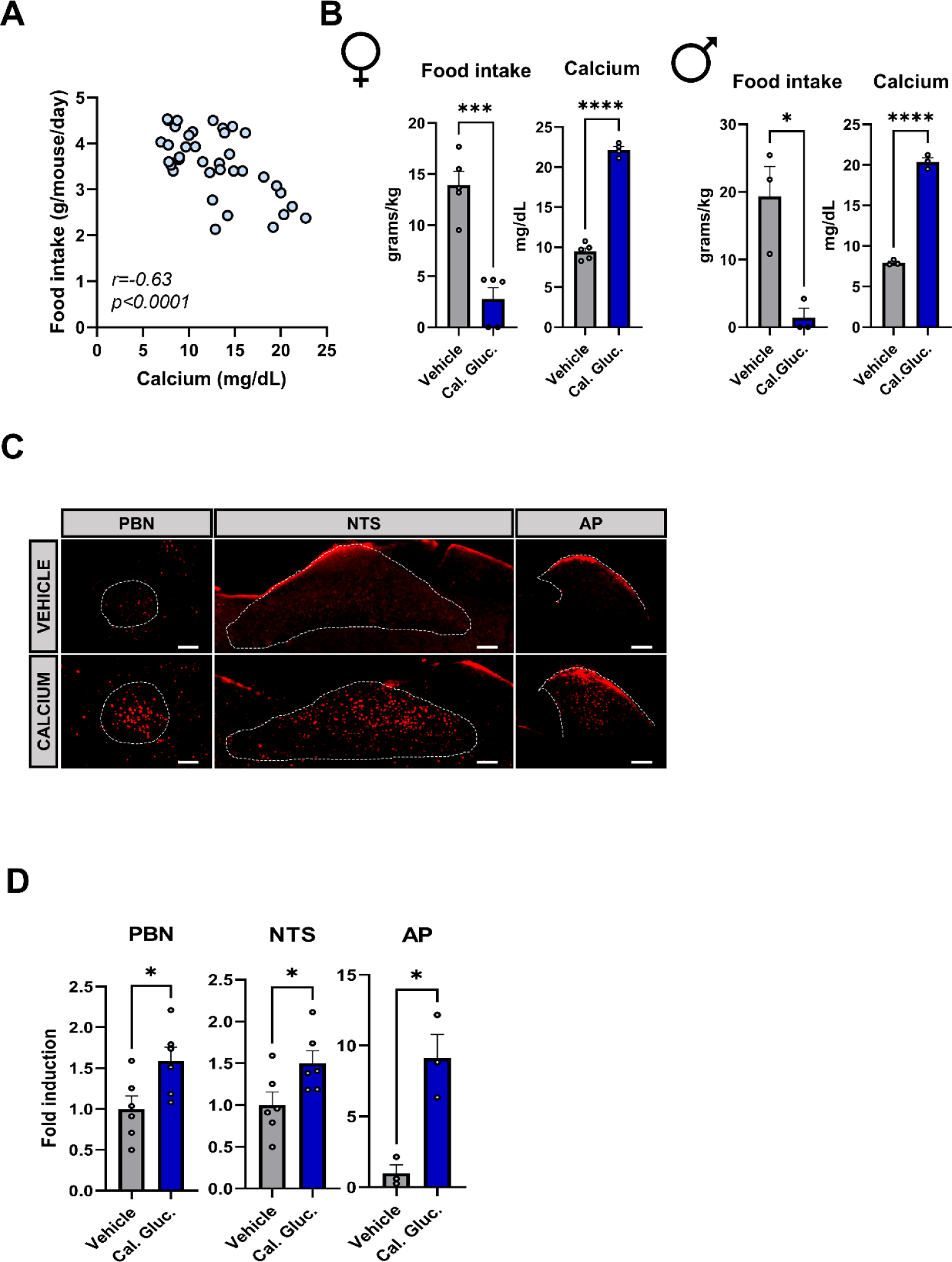
Inducing hypercalcemia in WT mice causes anorexia and activates neurons in the AP/NTS/PBN circuit. A) Pearson’s correlation analysis of the means of daily food intake and serum calcium levels (Pearson’s r = −0.63, p <0.0001) was calculated from the results obtained in the experiments from Figures 3, 4 & 5. B) Food intake after an overnight fast and serum calcium levels in 10-week-old female and male FVB WT mice injected i.p. with 1 mmol/kg calcium gluconate or vehicle (saline) every 30 min for 2h. Calcium levels were measured at the end of the 2-hour interval. C) Representative images of c-Fos (red fluorescence) immunostaining of vibratome sections of the AP, NTS and PBN in the sagittal plane. D) Quantification of c-Fos positive cells per unit area relative to control treated with vehicle. Scale bar: 100µm. Bars represent mean ± SEM, a minimum n=3. ****p<0.0001 ***p<0.001 *p<0.05.

Hypercalcemia has long been described to cause anorexia, nausea and vomiting in humans. Therefore, we hypothesized that circulating calcium released from the skeleton might be directly responsible for reducing food intake in Tet-PTHrP,PyMT mice on Dox. In support of this hypothesis, we found a significant negative correlation between circulating calcium levels and food intake across multiple different experiments in Tet-PTHrP;PyMT mice on Dox or controls (Fig. 5A). We next tested whether hypercalcemia in WT mice without tumors would impact food intake. We found that 4 injections of IP calcium gluconate over 2h raised circulating calcium levels in FVB mice to the same levels that we observed in Tet-PTHrP,PyMT mice on Dox ^44^ (Fig. 5B). Interestingly, inducing acute hypercalcemia in normal female mice caused an 80% reduction in food intake, compared to controls injected with vehicle (Fig. 5B). Moreover, brain sections of mice injected with calcium gluconate showed a marked increase in c-Fos expression in the AP, NTS and PBN, demonstrating that hypercalcemia, itself, can activate this hindbrain circuit and suppress food intake (Fig. 5C&D). Finally, calcium gluconate injections in male mice also caused hypercalcemia which led to a 90% reduction in food intake over the course of the experiment, demonstrating that hypercalcemia-induced-anorexia is conserved in both sexes (Fig. 5B).

### Hypercalcemia activates CaSR^AP^ neurons

Results from recent single nucleus RNA sequencing (snRNAseq) studies have found that *Casr* mRNA is expressed in glutamatergic neurons in the AP ^21,22^. Similarly, we interrogated the Allen Mouse Brain Atlas and noted that there are high levels of *Casr* mRNA detected in the AP from C57BL/6j mice, but barely detectable levels of *Casr* mRNA expression in the NTS, PBN, PVN or AN (Fig. 6A)^45^. We validated these findings by performing immunofluorescence staining with a CaSR antibody and by *in situ* hybridization (ISH) using the commercial, RNAscope platform. *Casr* mRNA was expressed in the AP but not in the NTS or PBN (Extended data Fig. 6A). Likewise, immunofluorescence for CaSR protein was evident within the AP (Extended data Fig. 6B).

**Figure 6.**
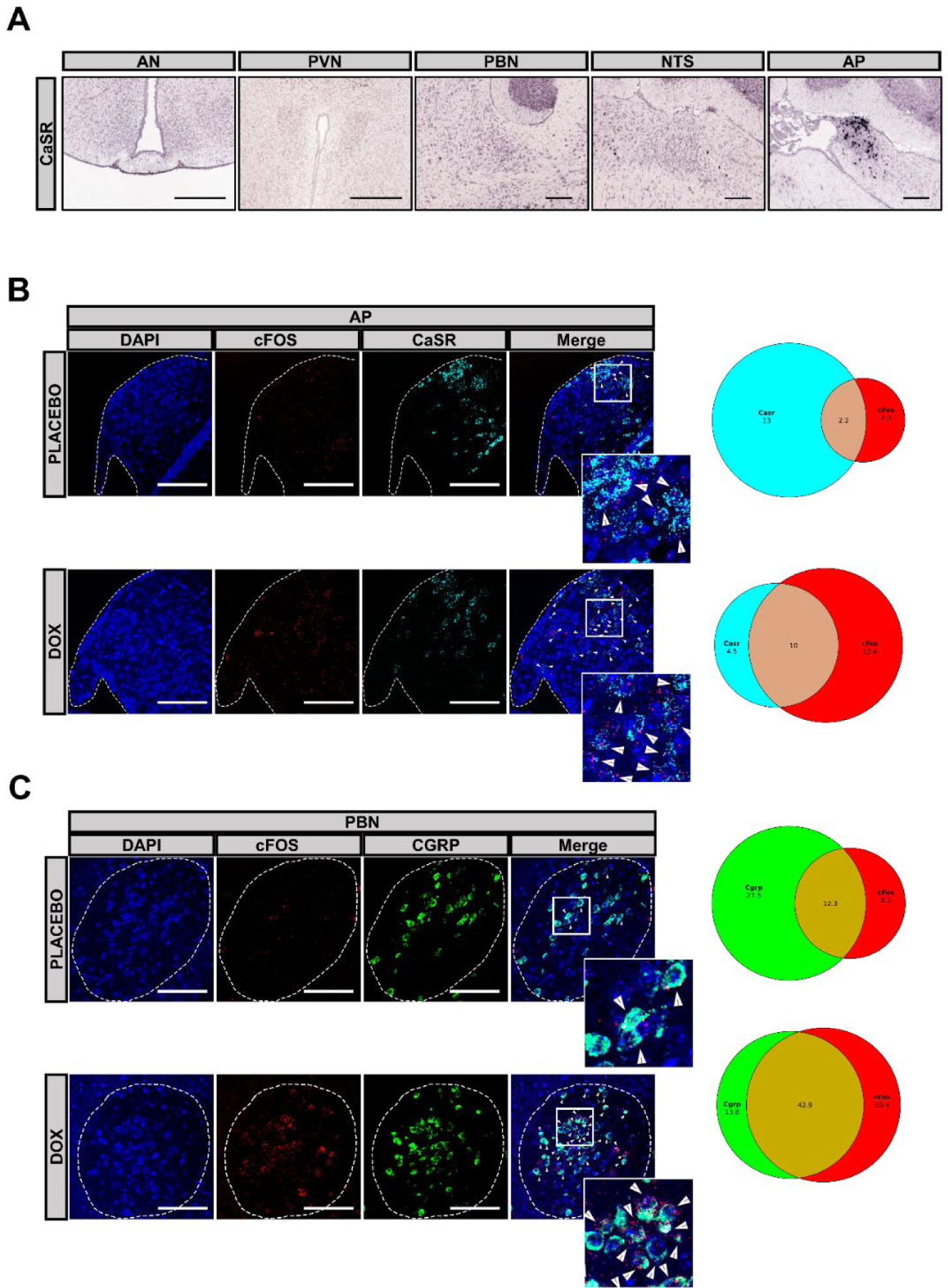
Tumor overexpression of PTHrP activates CaSR^AP^ and CGRP^PBN^ neurons. A) RNA *in situ* hybridization data from the Allen Brain Atlas showing strong expression of *Casr* mRNA in the Area Postrema (AP) and little or no *Casr* mRNA expression in the Arcuate Nucleus (AN), Paraventricular Nucleus (PVN), Parabrachial Nucleus (PBN), and Nucleus Tractus Solitarius (NTS). Scale bars: 210µm. B) Two-color fluorescent *in situ* hybridization of *Cfos* mRNA (red) and *Casr* mRNA (cyan) in the AP from sagittal sections of Tet-PTHrP;PyMT mice on placebo or Dox. C) *Cfos* mRNA (red) and *Cgrp* mRNA (green) in the PBN from sagittal sections of Tet-PTHrP;PyMT mice on placebo or Dox; Scale bars: 100µm. Insets showed magnified region as indicated by the white box outlined on the image. The numbers of individual and co-labeled cells from two separate mice were counted, pooled and expressed as the percentage of total nuclei in the pie graphs shown to the right of the corresponding images.

We next examined whether PTHrP-induced hypercalcemia specifically activates CaSR-expressing neurons within the AP (CaSR^AP^) by performing RNAscope ISH using probes against *Casr* and *cfos* on brain sections from Tet-PTHrP;PyMT mice treated with Dox or Placebo (Fig. 6B). The number of nuclei counted in the AP from both groups was similar (929 in Tet-PTHrP;PyMT mice on Dox vs 936 in Tet-PTHrP;PyMT on Placebo) and so was the percentage of cells expressing *Casr* (14.5% and 15.2% of total cells respectively). The percentage of *Cfos*+ cells was 5 times higher in the Dox treated animals compared to placebo-treated animals. Importantly, the percentage of cells expressing both *Casr* and *Cfos* mRNA was 4.5-fold higher with Dox treatment. The percentage of *Cfos*+ cells in Dox-treated animals was also increased by 3-fold in the NTS (Extended data Fig. 6C) and in the PBN (Fig. 6C). These ISH results are consistent with the previous results demonstrating that PTHrP overexpression increased c-Fos immunofluorescence in the AP, NTS and PBN. Previous reports have shown that signals from the AP and NTS activate neurons in the PBN expressing calcitonin gene-related peptide (CGRP^PBN^), and that these CGRP^PBN^ neurons are critical for inhibiting feeding ^23,46^. Therefore, we also examined *Cgrp* mRNA expression in the PBN and, as expected, found that it is highly expressed in the PBN of both Tet-PTHrP;PyMT mice on or off Dox. Furthermore, there were dramatic increases of *Cfos* mRNA in CGRP^PBN^ neurons in Dox-treated mice as compared to controls (42.9% vs 12.3% of total nuclei) (Fig. 6C). Taken together, these results demonstrate that hypercalcemia activates CaSR-expressing neurons in the AP as well as CGRP-positive neurons in the PBN known to trigger anorexia.

To test whether hypercalcemia alone can activate CaSR^AP^ neurons, we performed ISH analysis in brains from WT animals treated with calcium gluconate or control (Extended data Fig. 7A). Again, the percentage of total cells expressing *Casr* in the AP was similar between groups (12.4% and 14.6%, respectively), but we found that calcium gluconate treatment caused a 5.2-fold increase in *Cfos* mRNA expression and a 11-fold increase in the percentage of *Casr*+/*Cfos*+ cells compared to controls. Overall, these results suggest that hypercalcemia activates CaSR^AP^ neurons, which, in turn, activate downstream CGRP^PBN^ neurons that do not express the CaSR.

**Figure 7.**
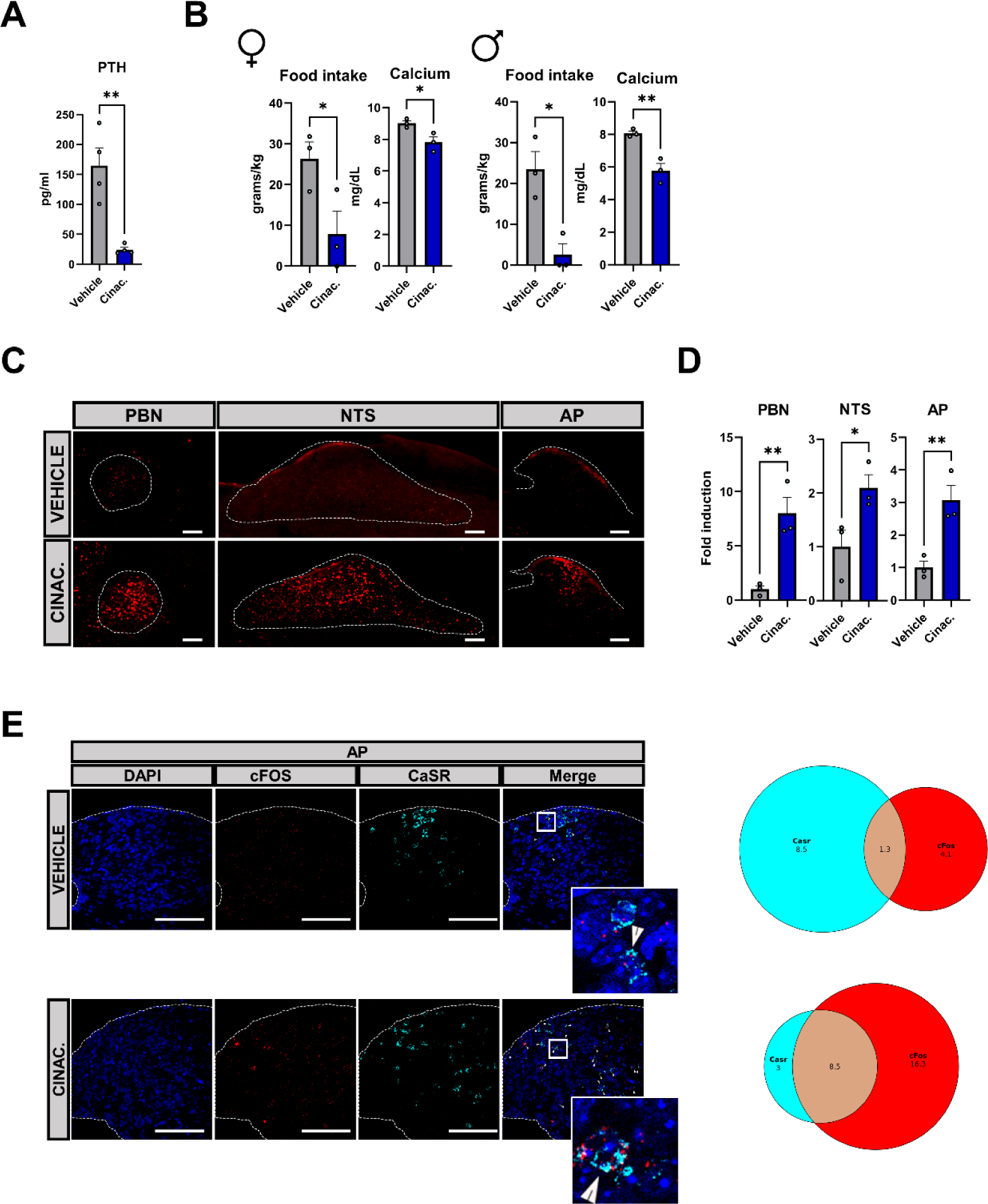
CaSR activation causes anorexia. A) Plasma PTH from 10-week-old WT, FVB mice injected with a single dose of 30 µmol/kg cinacalcet or vehicle [20% (2-Hydroxypropyl)-β-cyclodextrin (HβC)]. B) Food intake and serum calcium levels in female and male cinacalcet-treated WT mice as compared to mice treated with vehicle control. C) Representative images of c-Fos (red fluorescence) immunostaining of vibratome sections of the AP, NTS and PBN in the sagittal plane of mice treated with vehicle (top row) or cinacalcet (bottom row). D) Quantification of c-Fos positive cells per unit area in cinacalcet-treated WT mice relative to control WT mice treated with vehicle. Scale bar: 100µm. Bars represent mean ± SEM, n=3. **p<0.01 *p<0.05. E) Representative images of two-color fluorescent *in situ* hybridization of *Cfos* mRNA (red) and *Casr* mRNA (cyan) in the AP from sagittal sections of FVB mice treated with cinacalcet or vehicle; Scale bar: 100µm. The numbers of individual and co-labeled cells from two separate mice were counted, pooled and expressed as the percentage of total nuclei in the pie graphs shown to the right of the corresponding images.

### The CaSR Mediates the Effects of Calcium on the AP

To further test whether activation of the CaSR would stimulate the AP/NTS/PBN circuit and inhibit feeding behavior, we treated WT FVB mice with cinacalcet, an allosteric activator of the CaSR ^47^. Cinacalcet reduced circulating calcium and suppressed PTH levels, confirming its ability to activate the CaSR in the parathyroid glands (Fig. 7A&B). As compared to mice treated with vehicle, cinacalcet treatment also reduced food intake by 70% in females and 90% in males, mimicking the effects of high calcium even in the context of reduced circulating calcium levels (Fig. 7B). Consistent with the suppression of food intake, cinacalcet also increased nuclear C-Fos immunofluorescence in the AP, NTS and PBN of female mice (Fig. 7C&D). We also examined C-fos mRNA expression in CaSR-positive neurons in the AP in response to cinacalcet. Upon cinacalcet treatment, the percentage of *Cfos* expressing cells increased by almost 5-fold in the AP and the number of *Casr*+/*Cfos*+ cells in the AP increased by 6.3-fold compared to controls (Fig. 7E).

### Cinacalcet inhibits food intake and causes weight loss in HFD-obese mice

CaSR-expressing neurons in the AP also express the glucagon like peptide 1 receptor (GLP1R, Extended data Fig. 7B) ^21,22^. Semaglutide is a long-acting, GLP1R agonist that has been shown to inhibit food intake and cause weight loss in both pre-clinical rodent studies as well as in clinical trials in obese patients ^48–51^. Semaglutide may cause weight loss through several mechanisms but in mouse models of HFD-induced obesity, it activates GLP1R^AP^ neurons that connect to CGRP^PBN^ neurons ^23,28,52,53^. Therefore, we hypothesized that pharmacological activation of the CaSR might mimic the actions of GLP1R agonists to inhibit food intake and induce weight loss in diet-induced obese mice. To test this idea, we used a standard model of diet-induced obesity by feed C57/Bl6 male mice a HFD (45 Kcal % fat) for 12 weeks. Once mice were obese, we then treated them with daily injections of cinacalcet or vehicle for 7 days measuring daily changes in food intake and weight. Treatment with cinacalcet reduced food intake, with the maximal suppression occurring within the first 2 days. This led to a 12% reduction of cumulative food intake (Fig. 8A). Cinacalcet treatment also caused weight loss, displaying a 10% reduction in body weight (Fig. 8B). However, cinacalcet was dissolved in 20% (2-Hydroxypropyl)-β-cyclodextrin (HβC), which has previously been shown to cause weight loss ^54^. As a result, the vehicle-treated controls also demonstrated a 5% reduction in body weight.

**Figure 8.**
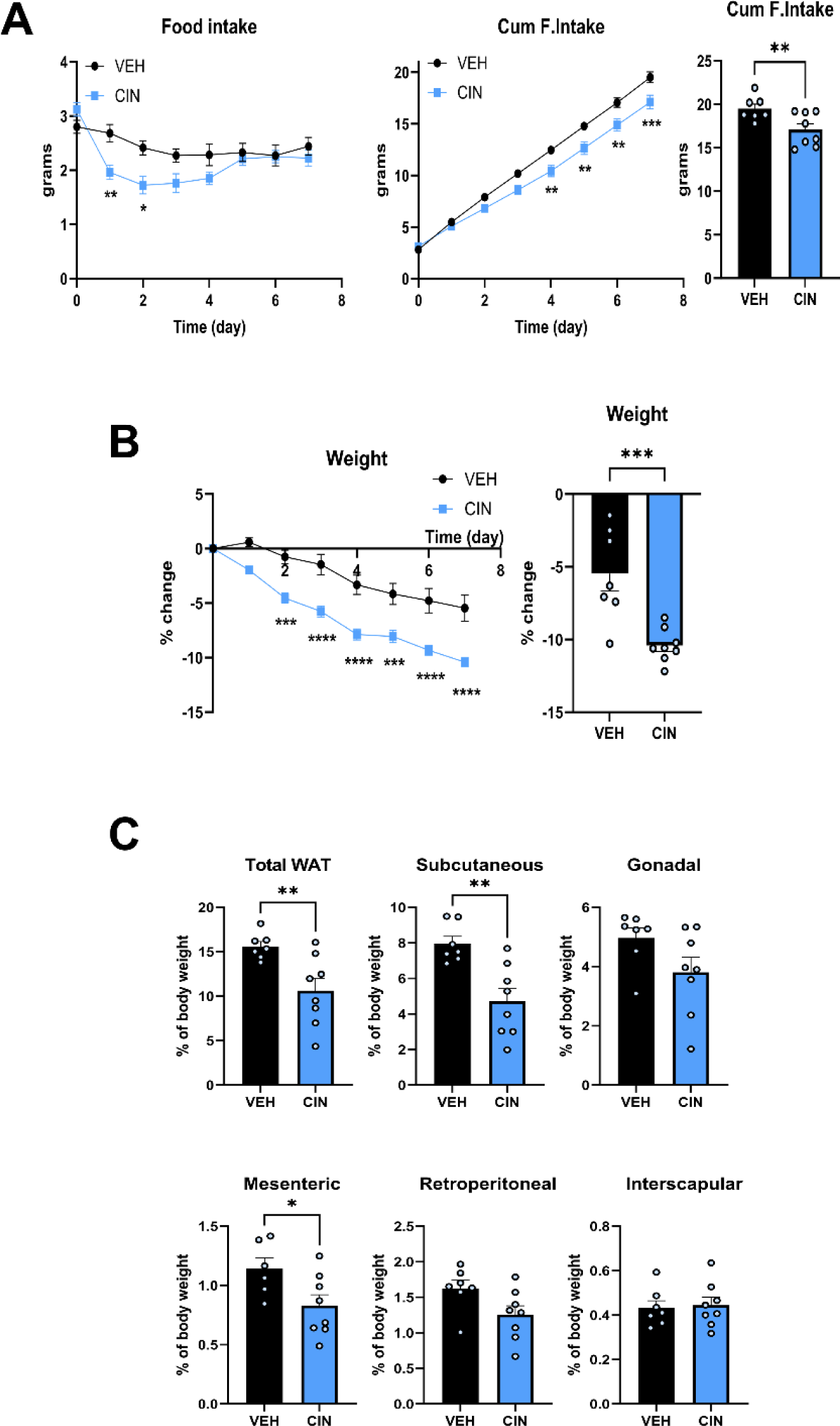
Cinacalcet treatment reduces food intake, body weight and adiposity in high-fat-fed mice. A) Food intake and cumulative food intake in cinacalcet-treated and vehicle-treated C57/Bl6 male mice fed with a high-fat diet (45 Kcal % fat) for 7 days. B) Weight change in cinacalcet-treated and vehicle-treated obese mice. C) Weight of different adipose tissue compartments from cinacalcet-treated and vehicle-treated obese mice at sacrifice. Weights are expressed as % of body weight. Bars represent mean ± SEM, n=8. ****p<0.0001 ***p<0.001 **p<0.01 *p<0.05.

At necropsy, fat depots from all mice were dissected and weighed. There was less total WAT in cinacalcet-treated HFD mice compared to the vehicle-treated controls (Fig. 8C), corresponding to a normalized total WAT loss of 30%. This was mainly due to marked reduction in adiposity within the subcutaneous and mesenteric WAT compartments (Fig. 8C).

These results in HFD obese mice suggest that activation of the AP/NTS/PBN pathway by the AP CaSR might be useful to induce anorexia and weight loss in obesity. Given that calcimimetics, which activate the CaSR are already FDA approved and used in patients, these existing agents might be repurposed or redesigned as anti-obesity drugs.

## DISCUSSION

This study demonstrates that cancer induced hypercalcemia causes anorexia by activating calcium sensing receptor-expressing neurons in the area postrema. In a mouse model of transgenic breast cancer, PTHrP overexpression by mammary tumor cells greatly increased circulating PTHrP levels and caused severe hypercalcemia, faithfully mimicking the common clinical syndrome of humoral hypercalcemia of malignancy or HHM. As often occurs in patients with HHM, these mice also experienced anorexia, rapid weight loss and fat wasting. Anorexia and weight loss were accompanied by activation of neurons in the AP, the NTS and the PBN, a hindbrain circuit well described to inhibit food intake. Our data demonstrate that anorexia was caused by elevated levels of calcium rather than elevated levels of PTHrP. Preventing hypercalcemia without affecting PTHrP and PTHR1 signaling blocked activation of the AP/NTS/PBN circuit and prevented anorexia in tumor-bearing mice. Furthermore, administering calcium or cinacalcet, an allosteric CaSR activator, caused anorexia and activated the AP/NTS/PBN circuit in WT mice. The CaSR is highly expressed in neurons in the AP, and we found that hypercalcemia specifically activated CaSR^AP^ neurons that were also positive for *GLP1R* mRNA. This, in turn, activated CaSR-negative neurons in the NTS and CGRP-positive neurons in the PBN that have previously been shown to inhibit food intake in response to activation of AP neurons ^55–57^. Finally, treating obese WT mice with cinacalcet reduced food intake and caused weight suggesting that the CaSR in the AP might be a pharmacologic target for obesity.

Cancer anorexia cachexia syndrome (CACS) affects up to 80% of all patients with cancer and contributes to 20% of all cancer deaths ^7^. The pathophysiology of CACS is thought to involve the secretion of inflammatory cytokines, hormones, neuropeptides, neurotransmitters, and/or metabolites from tumors that, directly or indirectly, suppress food intake and cause muscle and fat catabolism leading to involuntary weight loss ^58,59^. Previous studies suggested that PTHrP could contribute to CACS in both mice and humans. Clinical studies have reported that circulating PTHrP levels predict weight loss in cancer patients ^11^. A series of reports utilizing xenografts of human PTHrP-secreting tumors in nude mice or peripheral PTHrP administration in WT mice also suggested that PTHrP caused weight loss by inducing anorexia in rodents ^9,10,60–63^. More recently, Kir et al. suggested that PTHrP caused weight loss and fat wasting by inducing adipocyte “browning” and a hypermetabolic state without changing food intake ^8,37^. Our data support that PTHrP overexpression by tumors is associated with severe anorexia, but we found no evidence for increased metabolic rate or adipocyte browning in our breast cancer model. Furthermore, unlike the studies of Hashimoto et al. and Suzuki et al., we found no consistent changes in the pattern of expression of classic orexigenic or anorexigenic peptides in the hypothalamus ^9,64^. Instead, we found a striking activation of the AP, the NTS and the PBN, which together constitute a well described hindbrain neural pathway regulating food intake ^21,23–26^. Finally, our data suggest that PTHrP does not cause anorexia through direct actions on the brain but by inducing bone resorption and producing systemic hypercalcemia. Calcium, in turn, can activate the CaSR located on a subset of AP neurons which have also been shown to be activated by GLP-1, growth differentiation factor 15 (GDF15) and amylin, all of which have been shown to induce anorexia, at least in part, through activating the same AP/NTS/PBN neural pathway ^22,28–30,56,57^. We do not fully understand the discrepancies between our findings and those of other investigators regarding the contribution of PTHrP to CACS. However, our inducible transgenic model led to rapid elevations of circulating PTHrP and calcium levels while the previous xenograft models led to much longer periods of PTHrP exposure and more gradual increases in calcium levels. Therefore, our results do not exclude the possibility that PTHrP has direct effects on the hypothalamus or adipocytes over a longer time scale. In fact, we demonstrate that activation of the PTHR1 receptor on cultured adipocytes does stimulate lipolysis, although not “browning”.

Our findings are consistent with recent studies that have also reported the expression of the CaSR on neurons in the AP. Two studies used single nucleus RNAseq and single nucleus ATACseq to characterize the different neuronal populations in the AP. Both demonstrated that the CaSR is expressed by a specific sub-group of glutaminergic neurons that also co-expressed GLP1R, GDNF family receptor α-like (GFRAL), calcitonin receptor (CALCR) and receptor activity modifying protein 3 (RAMP3), all of which have been implicated in the regulation of food intake and/or food aversion ^21,22^. These CaSR^AP^ neurons respond to the long-acting GLP1R agonist, semaglutide, and project to GLP1R-positive and/or CGRP-positive, but CaSR-negative neurons in the NTS and PBN. The AP/NTS/PBN circuit has previously been described to be critical for learning food avoidance and Zhang and colleagues demonstrated that cinacalcet administration activated GLP1R-positive neurons in the AP to reinforce a flavor avoidance response ^21^. Additionally, Huang et al. showed that injection of NPS R568 (an early calcimimetic) directly into the AP activated glutaminergic neurons that acutely inhibited food intake ^65^. We confirmed those observations by showing first, that the CaSR is expressed in the AP (and not in the NTS or PBN) and that the approximately 13% of AP cells that express the CaSR also express the GLP1R. Second, we show that hypercalcemia stimulates cFos expression in CaSR^AP^ and CGRP^PBN^ neurons, suggesting that high calcium levels activate the CaSR in the AP and ‘turn on’ this anorectic pathway to inhibit food intake. Finally, systemic cinacalcet administration stimulates CaSR^AP^ neurons and leads to inhibition of food intake, mimicking the effects of hypercalcemia. While hypercalcemia has long been described to cause anorexia, there has not been a clear molecular explanation for these findings. Our data as well as those of other investigators noted above now provide strong support that elevations in extracellular calcium are sensed by CaSR^AP^ neurons, which activate neurons in the NTS and PBN to cause food aversion and anorexia. Therefore, inhibition of this pathway might provide relief for patients with CACS, especially those with co-existing hypercalcemia.

Recent years have seen an increased focus on the hindbrain as an important area regulating appetite, and several FDA-approved and investigational weight loss drugs activate neurons within the AP, NTS and/or PBN to suppress appetite ^25^. Drugs, such as semaglutide, that target the GLP1R have been shown to be highly effective at inducing weight loss in obese patients principally by reducing appetite and food intake ^48–50^. In rodent models of diet-induced obesity, semaglutide was shown to activate GLP1R^AP^ and CGRP^PBN^ neurons ^23,28,52,53^. Here, we show that cinacalcet also caused a reduction in food intake and weight loss in diet-induced obese male mice. Further studies will be needed to evaluate the efficacy of different doses and routes of administration for cinacalcet, but these initial studies suggest that this already FDA-approved medication could be repurposed as a weight loss medication. The most recent trials in human patients have demonstrated increased efficacy for weight loss using compounds that activate both the GLP1R and the GIPR, which is also expressed in the AP ^66^. Therefore, it will also be interesting to determine whether cinacalcet could magnify the effects of GLP1R, GIPR and/or amylin receptor agonists. This might allow the use of calcimimetics to lower doses of other weight loss drugs to avoid nausea, a common side effect of these drugs also likely to be related to activation of the AP/NTS/PBN pathway ^51,67^.

In summary, we demonstrate that systemic hypercalcemia caused by tumor production of PTHrP is associated with rapid weight loss due to severe anorexia. Reductions in food intake correlate with the degree of hypercalcemia and are associated with activation of CaSR-expressing neurons in the AP that may also be targets of GLP1R agonists used for weight loss. Calcimimetics, already in clinical use, reproduce these effects and cause weight loss in diet-induced obese mice. Further investigation of the actions of the CaSR in the area postrema may prove useful for developing new therapies targeting CACS and/or obesity.

## EXTENDED DATA

**Extended Data Figure 1.**
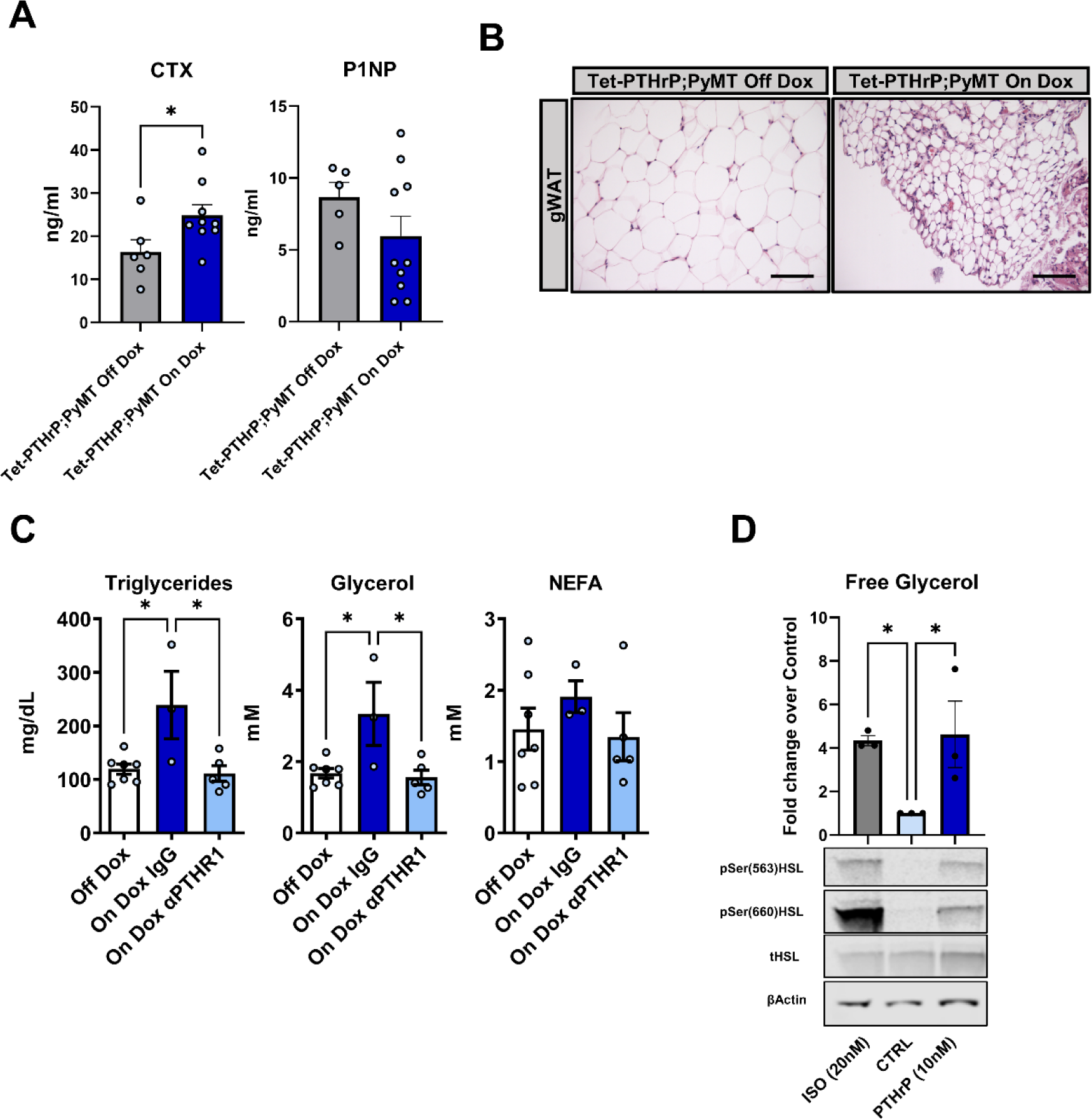
PTHrP increases bone resorption and induces lipolysis. A) 10-week-old Female Tet-PTHrP;PyMT mice with tumors were treated with Dox or vehicle for 4 days. A) Plasma CTX-1 and P1NP. B) Representative H&E images of gWAT from Tet-PTHrP;PyMT Off and On Dox. Scale bar: 100µm. C) Plasma triglycerides (TG), glycerol, and non-esterified fatty acids (NEFA) from 10-week-old, female Tet-PTHrP;PyMT mice on Dox treated with an anti-PTHR1 antibody or isotype control given by i.p. injection. Tet-PTHrP;PyMT mice off Dox were used as additional controls. D) Primary adipocyte cultures isolated from WT mice were stimulated with 10nM PTHrP (1-34) or 20 nM isoproterenol (ISO) for 30 min. Untreated cells (CTRL) were used for comparison. Lipolysis was measured as the glycerol concentration in the media. Aliquots of whole cell lysates were subjected to Western blot analysis using antibodies recognizing HSL, phosphorylated at S563 and S660. Antibodies against total HSL and β-Actin were used as controls. Bars in all graphs represent mean ± SEM. For each experiment, a minimum of 3 samples were analyzed. *p < 0.05.

**Extended data Figure 2.**
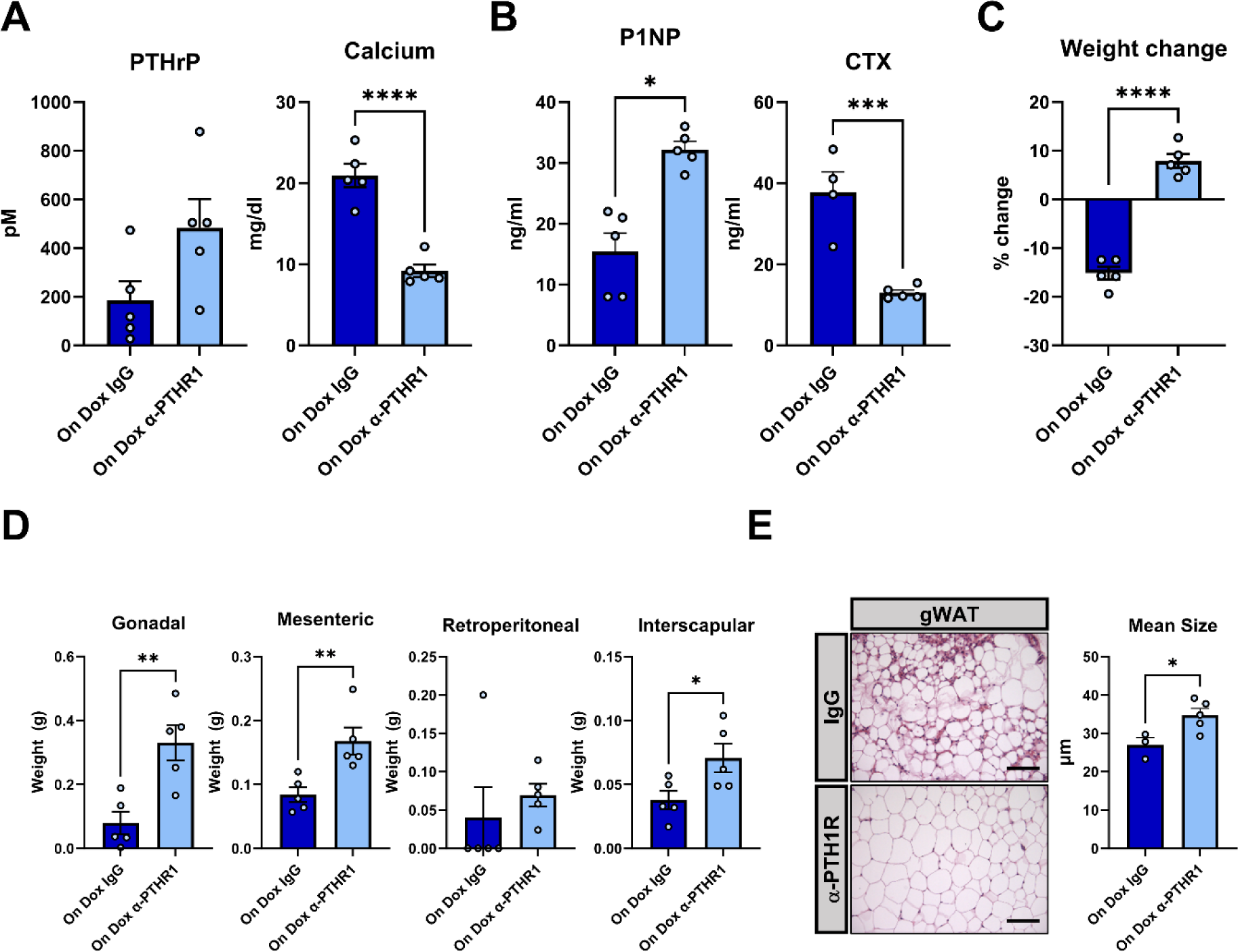
Blocking PTHR1 signaling corrects hypercalcemia and weight loss in tumor bearing mice. 10-week-old female Tet-PTHrP;PyMT mice on Dox were treated with an anti-PTHR1 antibody (αPTHR1) or isotoype control (IgG) given i.p. (A) Circulating plasma PTHrP and serum calcium concentration. (B) Plasma P1NP and CTX-1 levels. (C) Percent change in body weight over the course of the experiment. (D) Weights of different fat depots at time of sacrifice. (E) Representative H&E-stained histologic images of gonadal WAT from Tet-PTHrP;PyMT mice On Dox that were treated with either αPTHR1 or control IgG. Graph to the right represents the histomorphometric quantification of adipocyte size. Scale bars: 100µm. Bars in each graph represent mean ± SEM, n=5. ****p<0.0001 ***p<0.001 **p<0.01 *p<0.05.

**Extended Data Figure 3.**
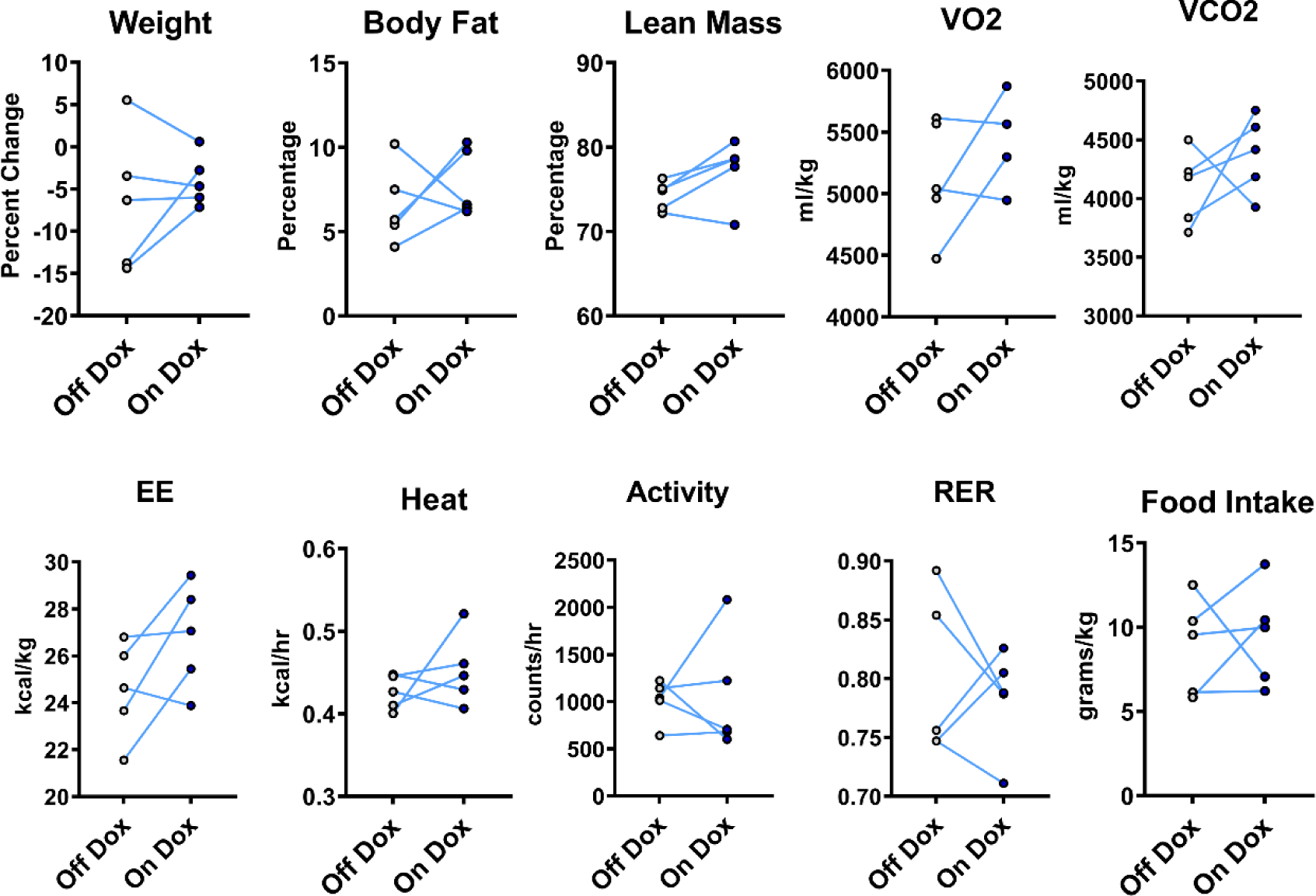
Doxycycline does not affect metabolism. Female, WT FVB mice were placed in Metabolic Cages and studied first on sucrose water for 5 days and then on dox for another 5 days. Weights and body compositions measured by ^1^H-NMR were obtained before and after each experimental period. Each pair of dots connected by a line represent data from the same mouse gathered during the 2 successive experimental periods; n=5. VO2, O2 consumption; VCO2, CO2 production; EE, energy expenditure; RER, respiratory exchange ratio. ****p<0.0001 ***p<0.001 **p<0.01 *p<0.05.

**Extended Data Figure 4.**
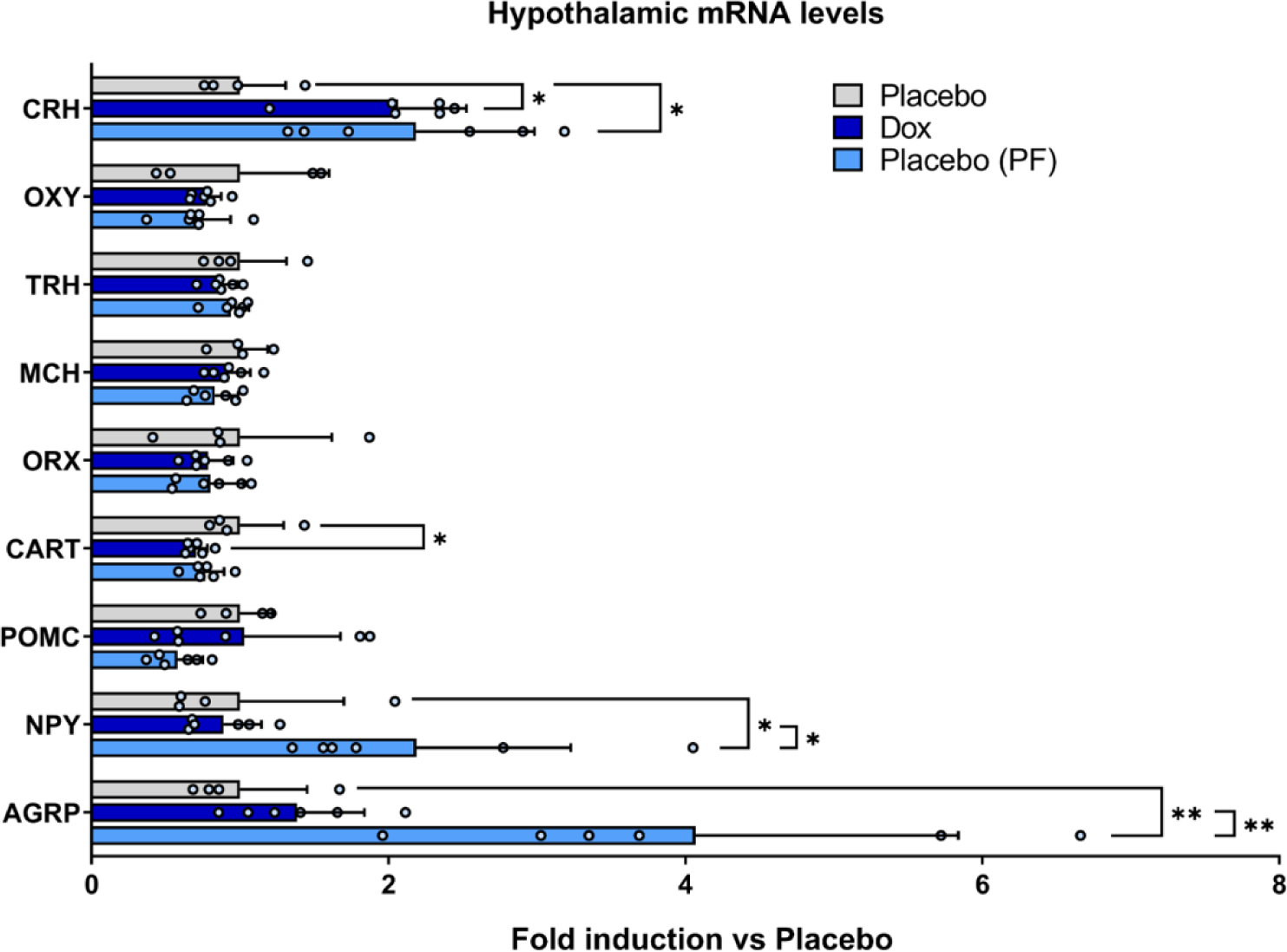
PTHrP does not modulate appetite regulating neuropeptide expression within the hypothalamus. The mediobasal hypothalamus was collected at sacrifice from Placebo, Dox or pair fed Placebo (Placebo (PF)) mice and total RNA was prepared. The expression of orexigenic (*Agrp, Npy, Orx, Mch*) and anorexigenic (*Cart, Pomc, Crh, Trh, Oxy*) peptide mRNA was determined by Real-time qPCR. *Hprt1* was used as housekeeping and the results are expressed as fold induction vs placebo. Bars represent mean ± SEM, a minimum n=4 mice were studied for each group. *p<0.05, ** p<0.01.

**Extended Data Figure 5.**
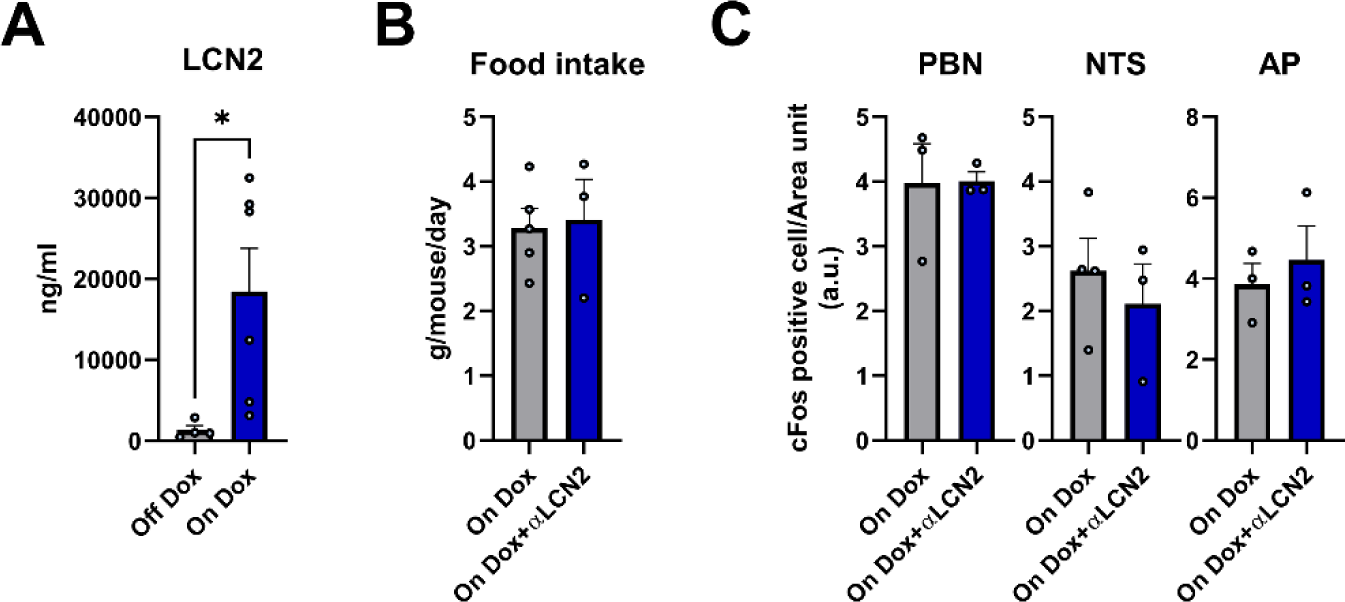
LCN2 does not mediate the PTHrP-induced anorexia. 10-week-old, female Tet-PTHrP;PyMT mice on Dox were treated with vehicle or an anti-LCN2 antibody for 4 days. A) Serum lipocalin-2 (LCN2) levels in Tet-PTHrP;PyMT mice on or off Dox. B) Daily food intake normalized by total body weight in Tet-PTHrP;PyMT mice on Dox treated with either anti-LCN2 antibody or control. C) Quantification of c-Fos positive cells per unit area from c-Fos immunostaining of vibratome sections of the AP, NTS and PBN. Bars represent mean ± SEM, at least 3 mice were studied in each group. *p<0.05.

**Extended Data Figure 6.**
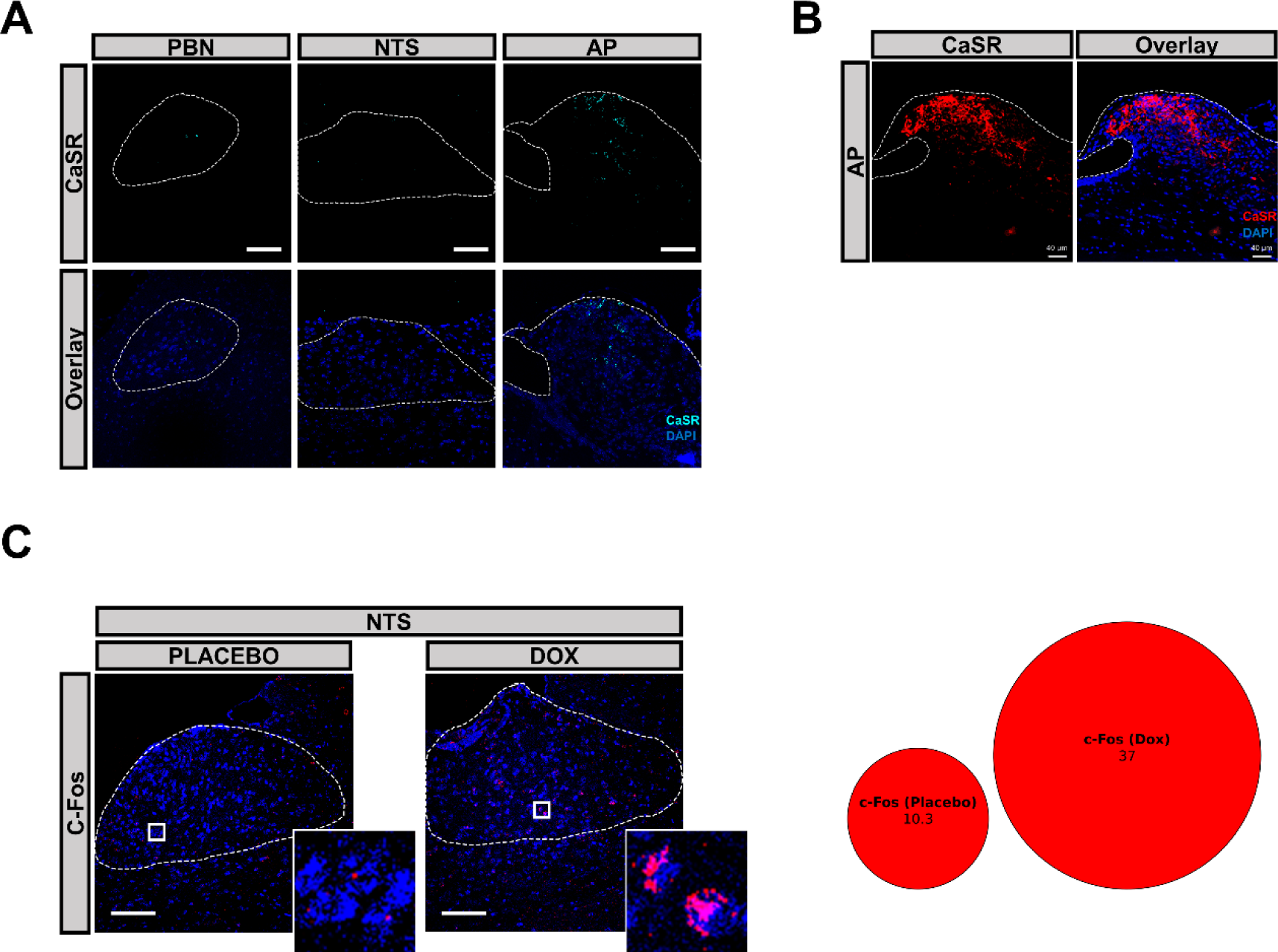
CaSR is expressed in the AP and its activation induces NTS c-Fos expression. A) Representative images of fluorescent *in situ* hybridization of *Casr* mRNA (cyan) in the AP, NTS and PBN from sagittal sections of FVB mice. B) Immunofluorescence staining for CaSR protein (red fluorescence) in AP sagittal sections of Tet-PTHrP;PyMT mice on Placebo. C) Representative images of fluorescent *in situ* hybridization analysis of *Cfos* mRNA (red fluorescence) in the NTS from sagittal sections of 10-week-old, Tet-PTHrP;PyMT mice on placebo vs. on Dox. The numbers of individual-labeled cells from two separate mice were counted, pooled and expressed as the percentage of total nuclei in the pie graphs shown to the right of the corresponding images. Scale bars: 100µm

**Extended Data Figure 7.**
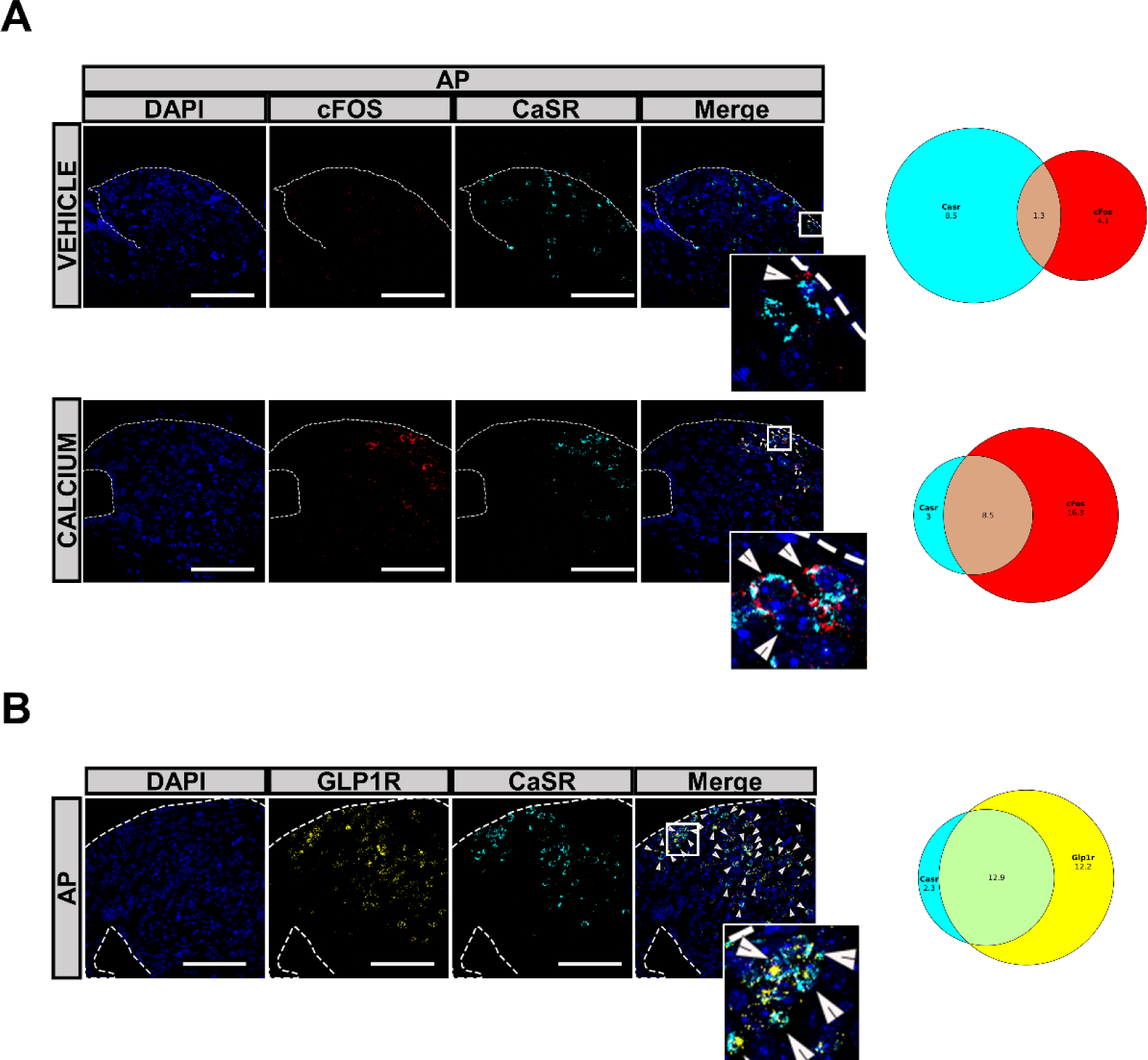
CaSR expressing neurons in the AP are activated by calcium and also express GLP1R. A) Two-color fluorescent *in situ* hybridization analysis of *Cfos* mRNA (red) and *Casr* mRNA (cyan) in the AP performed on representative sections from WT mice treated with vehicle (top row) or calcium gluconate (bottom row). The numbers of individual and co-labeled cells from two mice were counted, pooled and expressed as percentage of total nuclei (right). B) *Glp1r* mRNA (yellow) and *Casr* mRNA (cyan) in the AP of vehicle treated FVB mice. The numbers of individual and co-labeled cells from two mice were counted, pooled and expressed as the percentage of total nuclei (right). Scale bars: 100µm.

